# Recurrent neural network models of multi-area computation underlying decision-making

**DOI:** 10.1101/798553

**Authors:** Michael Kleinman, Chandramouli Chandrasekaran, Jonathan C. Kao

## Abstract

Cognition emerges from coordinated computations across multiple brain areas. However, elucidating these computations within and across brain regions is challenging because intra- and inter-area connectivity are typically unknown. To study coordinated computation, we trained multi-area recurrent neural networks (RNNs) to discriminate the dominant color of a checker-board and output decision variables reflecting a direction decision, a task previously used to investigate decision-related dynamics in dorsal premotor cortex (PMd) of monkeys. We found that multi-area RNNs, trained with neurophysiological connectivity constraints and Dale’s law, recapitulated decision-related dynamics observed in PMd. The RNN solved this task by a dynamical mechanism where the direction decision was computed and outputted, via precisely oriented dynamics, on an axis that was nearly orthogonal to checkerboard color inputs. This orthogonal direction information was preferentially propagated through alignment with inter-area connections; in contrast, color information was filtered. These results suggest that cortex uses modular computation to generate minimal sufficient representations of task information. Finally, we used multi-area RNNs to produce experimentally testable hypotheses for computations that occur within and across multiple brain areas, enabling new insights into distributed computation in neural systems.

## Introduction

Decision-making, multisensory integration, attention, motor control, and timing emerge from the coordination of multiple interconnected brain areas (Szczepanski et al., 2010, Kalaska & Crammond, 1992, Remington et al., 2018, Yamagata et al., 2012, Yau et al., 2015, Cisek, 2012, Pinto et al., 2019, Siegel et al., 2015, Freedman & Assad, 2016). While neural activity in a particular area can contain behaviorally relevant signals, such as choices or percepts, it is often unclear if these signals originate within the area or are inherited from upstream brain areas (Cisek, 2012). Understanding the neural bases of these behaviors necessitates an understanding of the intra- and inter-area dynamics, that is, how neural activity evolves within and between brain areas (Brown et al., 2004, Gerstein & Perkel, 1969, Ghazanfar et al., 2008, Buschman & Miller, 2007, Saalmann et al., 2007, Gregoriou et al., 2009, Semedo et al., 2019, Tauste Campo et al., 2015). However, we currently lack clear hypotheses for how distinct brain-area dynamics and connectivity relate to computation. To address this gap, we use multi-area recurrent neural networks (RNNs) to model, probe, and gain insight into how behaviorally relevant computations emerge within and across multiple brain areas.

Optimized feedforward and recurrent neural networks are emerging tools to model computations associated with visual (Yamins et al., 2014, Yamins & DiCarlo, 2016), cognitive (Mante et al., 2013, Song et al., 2016, Rajan et al., 2016, Chaisangmongkon et al., 2017, Song et al., 2017, Yang et al., 2019, Orhan & Ma, 2019), timing (Laje & Buonomano, 2013, Goudar & Buonomano, 2018, Remington et al., 2018), navigation Banino et al. (2018), and motor tasks (Hennequin et al., 2014, Sussillo et al., 2015, Michaels et al., 2016, Stroud et al., 2018). RNNs transform experimenter-designed task inputs into behavior-related outputs through recurrent dynamics. Its artificial units often exhibit heterogeneous responses and population dynamics observed in brain areas implicated in cognitive and motor tasks (Barak, 2017, Mante et al., 2013, Hennequin et al., 2014, Sussillo et al., 2015, Song et al., 2016, 2017). If artificial units resemble cortical neurons, RNNs are subsequently analyzed to propose hypotheses for how a local computation occurs in a brain area (Mante et al., 2013, Sussillo et al., 2015, Chaisangmongkon et al., 2017, Remington et al., 2018). An important advantage of RNNs is that the activity of all artificial units and their recurrent connectivity are fully observed. It is therefore possible to engineer (Mastrogiuseppe & Ostojic, 2018) and reverse engineer (Sussillo & Barak, 2013) RNNs by analyzing their activity and recurrent connectivity, providing mechanistic insight into cortical computation (Mante et al., 2013, Chaisangmongkon et al., 2017, Remington et al., 2018).

Traditionally, RNNs have provided insight into local computations, however, there has been limited insight into multi-area computation (Song et al., 2016, Pinto et al., 2019, Michaels et al., 2019). To study multi-area computation, we explicitly constrained RNNs to have multiple recurrent areas, which we refer to as multi-area RNNs. We used these multi-area RNNs to study decision-making, a cognitive process known to involve multiple areas including the prefrontal, parietal, and premotor cortex (Mante et al., 2013, Chandrasekaran et al., 2017, Kaufman et al., 2015, Coallier et al., 2015, Pinto et al., 2019, Gold & Shadlen, 2007, Freedman & Assad, 2016, Romo & de Lafuente, 2013). Multi-area RNNs enable us to investigate several questions. Most broadly, what are the roles of within-area dynamics and inter-area connections in mediating distributed computations? What role do inter-area feedforward and feedback connections play in propagating information and rejecting noise? How do intra-area dynamics and inter-area connections coordinate to solve a task?

We use multi-area RNNs to study these questions in a decision-making task where premotor cortex and upstream areas are known to perform distinct computations. In our “Checkerboard Task” (Chandrasekaran et al., 2017, Coallier et al., 2015, Wang et al., 2019), shown in Fig. 1a), a monkey was first shown red and green targets whose location (left or right) was randomized on each trial (‘Targets’ epoch in Fig. 1a). The monkey was subsequently shown a central static checkerboard composed of red and green squares. The monkey was trained to discriminate the dominant color of the static checkerboard and reach to the target matching the dominant color. Since the target locations were randomized on each trial, this task separates the reach direction decision from the color decision (Freedman & Assad, 2011). Neurons in the caudal portion of dorsal premotor cortex (PMd) show strong covariation with the direction decision, but only very weak association with the color decision (Chandrasekaran et al., 2017, Wang et al., 2019, Coallier et al., 2015). This result suggests that the transformation from a color to direction decision is computed upstream of PMd, implicating multiple brain areas in solving this task.

**Figure 1:**
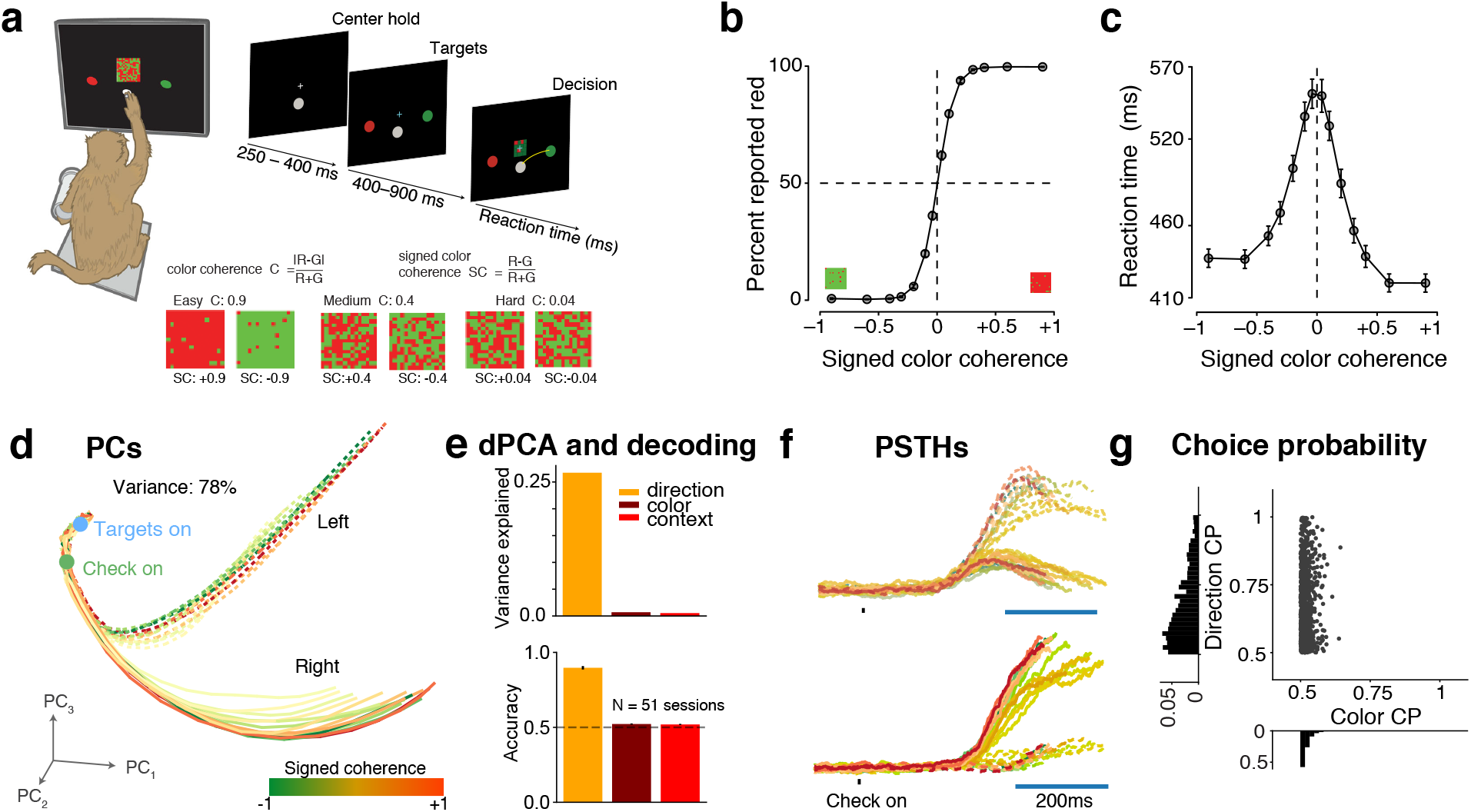
PMd data suggests multiple brain areas are implicated in a perceptual decision making task. **(a)** Perceptual decision-making task: A monkey, after holding a center location for 250 − 400 ms, is presented with a red and green target. The location of these targets (i.e., left or right) is randomly chosen on each trial. After a holding period of 400 − 900 ms, the monkey is presented a 15 × 15 checkerboard, composed of red and green squares. To successfully complete the trial, the monkey must reach to the target matching the dominant color of the checkerboard. **(b,c)** Psychometric curve (panel b) and reaction time (panel c) from one monkey from prior experimental work Chandrasekaran et al. (2017). Figure reproduced from (Chandrasekaran et al., 2017) with permission. **(d)** PMd neural trajectories in the top 3 PCs. Color reflects signed color coherence, with darker shades of red (green) indicating more red (green) checkerboards. Right (left) reaches are denoted by solid (dotted) lines. **(e)** (Top) Variance captured by dPCA axes for the color decision, target configuration (context), and direction decision. (Bottom) Decode accuracy of the direction decision, color decision, and context in PMd sessions with U-probes and multiple neurons. **(f)** Representative PMd PSTHs aligned to checkerboard onset. X-axes is time in ms, Y-axes are firing rate in Spks/s. PMd PSTHs primarily separate based on the direction, not the signed coherence. **(g)** Direction and color choice probability (CP) for all recorded PMd units.

We trained multi-area RNNs to perform the Checkerboard task. We found that, when incorporating Dale’s law and neurophysiological constraints into training, PMd-resembling dynamics emerged in multi-area RNNs. Specifically, the multi-area RNN’s output area (1) resembled PMd in single unit statistics and neural population activity, and (2) represented the direction decision but not the color decision. These RNNs employed a dynamical mechanism where the direction decision was computed and outputted on an axis that was nearly orthogonal to the checkerboard color inputs and color representations. Inter-area connections preferentially propagated this orthogonal direction information and attenuated color information. Together, our results demonstrate that multi-area computation involves coordinated intra-area dynamics that are selectively propagated to subsequent areas. Our models and analyses provide a framework for studying distributed computations involving multiple areas in neural systems.

## Results

### Decision-making involves multiple brain areas

We analyzed the activity of PMd neurons in monkeys performing the Checkerboard Task (Chandrasekaran et al., 2017, Fig. 1a). Monkeys discriminated the dominant color of a central static checkerboard (15 × 15 grid) composed of red and green squares. The number of red and green squares was randomized across trials, leading to different levels of discrimination difficulty. The signed color coherence linearly indicates the degree to which color is dominant in the checkerboard, with −1 corresponding to completely green, 0 to equal numbers of red and green squares, and +1 to completely red. If the monkey reached to the target matching the dominant checkerboard color, the trial was counted as a success and the monkey received a juice reward. Critically, the color decision (red or green) was decoupled from the direction decision (left or right) because the left and right target identities were random on every trial. Fig. 1b,c show the monkey’s psychometric and reaction time (RT) behavior on this task. Monkeys made more errors and reacted more slowly for more ambiguous checkerboards compared to the almost completely red or completely green checkerboards (Chandrasekaran et al., 2017).

In the Checkerboard Task, neural activity in PMd, an area associated with somatomotor decisions, principally reflects the direction decision (left or right) and has minimal representations associated with the dominant color of the checkerboard (red or green) (Chandrasekaran et al., 2017, Wang et al., 2019, Coallier et al., 2015). To summarize this phenomenon, we show the principal components (PCs) of the PMd neural population activity in Fig. 1d. These PC trajectories separated based on the eventual reach direction (right reaches in solid, left in dotted), not color (red and green overlapping). We identified principal axes via dPCA that maximized variance related to the target configuration (context), color decision, and direction decision (see Methods). The direction axes captured significant variance (26.7%) while the color and context axes captured minimal variance (0.7%, 0.5%, respectively), as shown in Fig. 1e. This means that there is no independent color and context variance in PMd. It is possible, however, that there is direction-dependent color variance that is averaged away during marginalization when computing the dPCA variance (Kobak et al., 2016). Given simultaneously recorded data, a more appropriate measure of representation is the decode accuracy of direction, color and context. Across sessions where we recorded multiple units from U-probes, the direction decision could be decoded from PMd activity significantly above chance (accuracy: 0.89, *p* < 0.01, bootstrap), but the color decision and context decode accuracy were not significantly above chance in any session (overall accuracies: 0.52 and 0.52, respectively, Fig. 1e).

Single neurons also had minimal color separation in individual PSTHs (e.g., Fig. 1f). To summarize this effect in single neurons, we computed the choice probabilities (CPs) reflecting how well the direction decision (direction CP) and color decision (color CP) could be decoded. PMd units generally had near chance color CP (0.5), but moderate to high direction CP, as shown in Fig. 1g. Together, these results demonstrate that PMd largely represents direction-related signals, but not the color decision or target configuration context.

Since PMd activity minimally represents the color of the checkerboard or the target configuation, we reasoned that checkerboard and target inputs are transformed into a direction signal upstream of PMd and that multiple brain areas are necessary for solving this task. Brain areas, including the dorsolateral prefrontal cortex (DLPFC), and the ventrolateral prefrontal cortex (VLPFC), have been implicated in related sensorimotor transformations (Wise & Murray, 2000, Hoshi, 2013, Yamagata et al., 2012, Badre et al., 2009, Freedman & Assad, 2016). VLPFC is part of the ventral pathway and is connected to the inferotemporal cortex (Gerbella et al., 2009, Yeterian et al., 2012), which processes color and object related information (Conway, 2009). Color-related inputs from the ventral pathway comes into PMd through a polysynaptic pathway via projections from the VLPFC to DLPFC (Takahara et al., 2012). Thus, these areas are anatomically connected in a loosely hierarchical manner: VLPFC is connected to DLPFC, which is connected to rostral PMd (PMdr) but not PMd, and PMdr sends strong projections to PMd (Barbas & Pandya, 1987, Luppino et al., 2003).

### PMd-like representations emerge in optimized multi-area RNNs

Given this anatomical and physiological evidence, we hypothesized that the last area of an optimized multi-area RNN would more closely resemble PMd, receiving transformed direction signals computed using the checkerboard coherence and target configuration from upstream areas. We trained multi-area RNNs to perform the Checkerboard Task. Training details, including input and output configuration, optimization, and architectural constraints, are detailed in the Methods. Briefly, the multi-area RNN processed the target context and checkerboard coherence to produce two decision variables that accumulated evidence for a left and right decision (Fig. 2a). The RNN had 3 areas, obeyed Dale’s law (Song et al., 2016), and had approximately 10% feedforward and 5% feedback connections between areas based on projections between prefrontal and premotor cortex in a macaque atlas (Markov et al., 2014). RNN psychometric and RT curves exhibited similar behavior to monkeys performing this task (Fig. 2b,c; across several RNNs, see Fig. S1).

**Figure 2:**
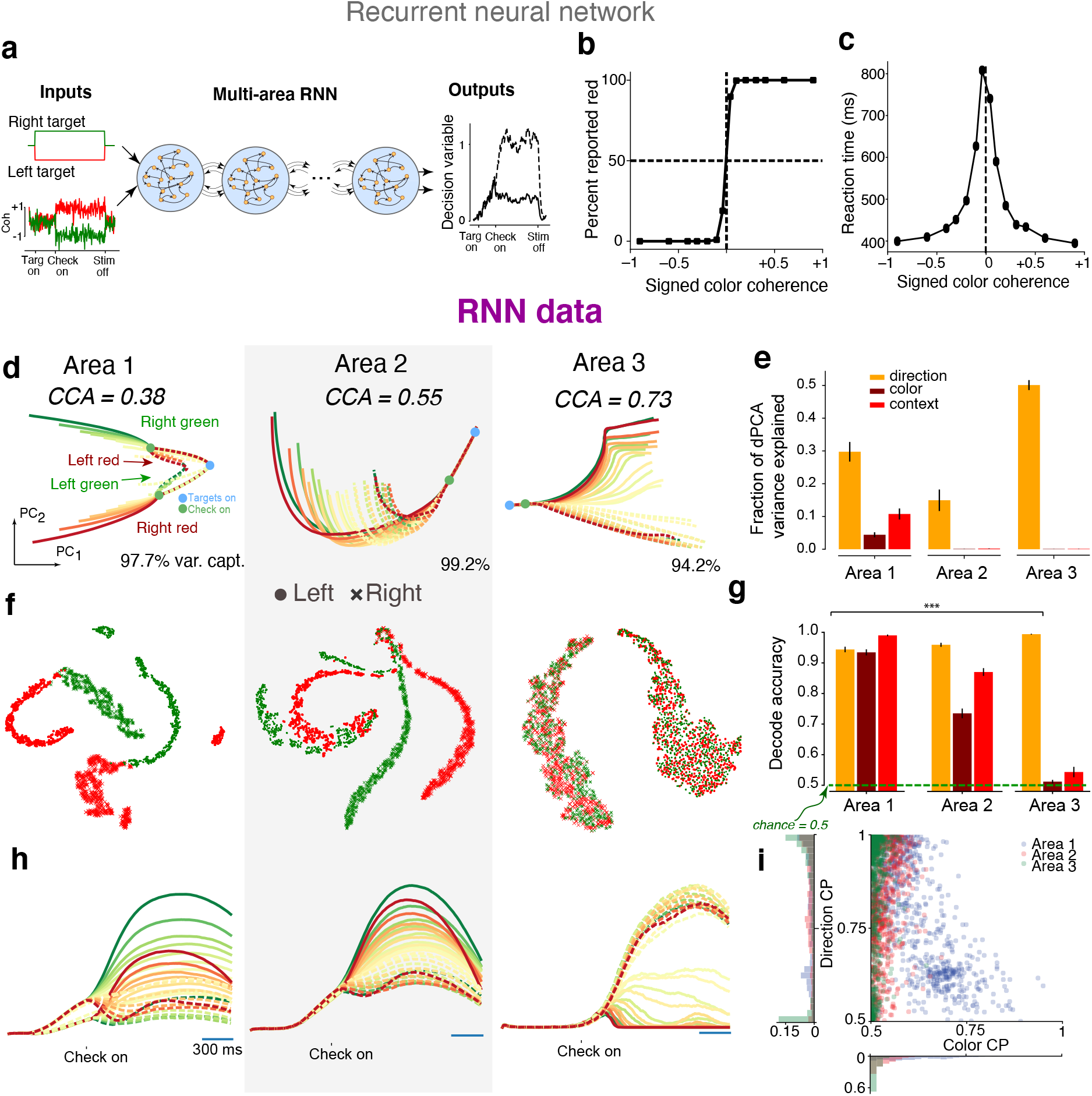
PMd-resembling dynamics emerge in optimized multi-area RNNs. **(a)** Multi-area RNN configuration. The RNN received 4 inputs. The first two inputs indicated the identity of the left and right targets, which was red or green. These inputs were noiseless. The last two inputs indicated the value of the signed color coherence (proportional to amount of red in checkerboard) and negative signed color coherence (proportional to amount of green in checkerboard). We added independent Gaussian noise to these signals (see Methods). The network outputted two analog decision variables indicating evidence towards the right target (solid line) or left target (dashed line). A decision was made in the direction of whichever decision variable passed a preset threshold (0.6) first. The time at which the decision variable passed the threshold was defined to be the reaction time. **(b,c)** Psychometric curve and reaction time for the multi-area RNN. **(d)** Neural trajectories in the top 2 principal components for each RNN area. **(e)** Variance captured by dPCA axes for color, context, and direction. Area 1 contains color, direction and context information, whereas Area 3 contains minimal variance for color and context. Note dPCA is a linear method that depends on marginalization. **(f)** Non-linear tSNE embedding of peri-movement activity in each area. Each dot is a trial, with red or green denoting the color decision and ‘.’ or ‘x’ denoting the direction decision. Unlike Areas 1 and 2, Area 3 only had two clusters separated based on the direction decision. **(g)** Decode accuracy of direction, color, and context in all three areas. Although Area 2 had little color and context variance, the color decision and context achieved significantly above chance decode accuracy (***, *p* < 0.01/9, corrected for multiple comparisons). Decoding the color decision from Area 3 activity achieved nearly chance decode accuracies (*p* = 0.05, one-tailed t-test). **(h)** Example PSTHs in each area. Note that PSTHs in Area 3 only separate by reach direction and not signed coherence, whereas in Areas 1 and 2, they separate by signed coherence and reach direction. **(i)** Choice probabilities for units in all areas (pooled over 8 RNNs).

The 3-area RNN had qualitatively different population trajectories across areas, shown in Fig. 2d. Area 1 had four distinct trajectory motifs corresponding to the four possible task outcomes (combinations of left vs right and red vs green decisions). PC_1_ primarily varied with direction, while PC_2_ varied with both the target context and red versus green checkerboards. In contrast, Area 2 and Area 3 population trajectories primarily separated on direction, not color, like in PMd. Area 3 trajectories most strongly resembled PMd trajectories (canonical correlations, *r* = 0.38, 0.55, 0.73 for Areas 1, 2, and 3; see Methods). We quantified the variance captured by dPCA principal axes for the context, color axis, and direction. We found that color axis variance decreased in later areas (Area 1: 5.6%, Area 2: 0.13%, Area 3: 0.07%, Fig. 2e). In contrast, Area 3 had the largest direction axis variance (Area 1: 30.9%, Area 2: 18.2%, Area 3: 48.5%, Fig. 2e). An important assumption of dPCA is that the neural activity can be decomposed as a sum of terms that depend solely on particular task variables (Latimer, 2019). The color variance found by dPCA indicate that color, on its own, did not account for a large fraction of the overall neural variance. However, it is possible there is significant color variance within a reach direction that dPCA, a linear dimensionality reduction technique, does not capture. When considering the direction dependent color variance, we found this was the case: color variance had larger values, but nevertheless decreased in later areas (Area 1: 81%, Area 2: 4.6%, Area 3: 0.5%).

As we are interested in whether the color information is contained is the representation, a more appealing measure is decode accuracy. If the color of the target can be decoded from the representation of neural activity, then color information is present in the representation. We performed nonlinear dimensionality reduction via t-distributed stochastic neighbor embedding (tSNE) (Maaten & Hinton, 2008), shown in Fig. 2f. These results suggest that Areas 1 and 2 contain color information, but Area 3 does not (color decisions overlap). We decoded the color decision and context (target configuration) from RNN units in each area (Fig. 2g, Area 1, 2, and 3 color accuracy: 0.93, 0.76, 0.51, and Area 1, 2, and 3 context accuracy: 0.99, 0.87, 0.54). Area 1 and 2 had above chance context and color decode accuracies (*p* < 0.01/9, 1-tailed t-test with Bonferroni correction), while Area 3 color and context decode accuracies were near chance, and most similar to PMd (Fig. 2g, color: *p* = 0.05, context: *p* = 0.024). The direction decision could be decoded significantly above chance in all areas (Fig. 2g, *p* < 0.01/9).

We also observed that Area 3 unit PSTHs more closely resembled PMd neuron PSTHs (e.g., Fig. 2h), and color CP progressively decreased in later areas (Fig. 2i). Area 3, like PMd, had many neurons with moderate to high direction CP, but low color CP. Together, this constellation of results shows that Area 3 of the multi-area RNN recapitulates key features of PMd activity, making this RNN a candidate model of decision-making in the Checkerboard Task.

We note that we also trained traditional single-area RNNs with Dale’s Law to perform this task. When trained with the task inputs described in Fig. 2a, we did not observe PMd-resembling dynamics emerge in these single-area RNNs (Fig. S2). Example principal components (Fig. S2c) and PSTHs (Fig. S2e) reveal an encoding of color, in contrast with PMd principal components and PSTHs. As we highlight in the Discussion, this result should not be interpreted as evidence that a single-area RNN cannot provide insights into PMd, but that for our task setup and specific inputs we chose, PMd-like dynamics did not naturally emerge through optimization across several hyperparameter settings (Fig. S5).

### Minimal color representations emerge in RNNs trained with neurophysiological constraints

Is our result an inevitable consequence of incorporating multiple areas? Stated differently, what design choices and hyperparameters lead to multi-area RNNs with minimal color representation? We varied RNN architectures and hyperparameters to determine the settings that led to the emergence of PMd-like minimal color representations. Critically, we found that the key design choices affecting our results were neurophysiological constraints. In contrast, our results were more robust to machine learning related hyperparameters. Unless otherwise noted, statistical tests are one-tailed t-tests across trained networks with Bonferroni correction for multiple comparisons.

#### Multiple areas

We first tested if minimal color representations were a simple consequence of multi-area computation. We trained 3-area RNNs that were “unconstrained” in that artificial units could have both excitatory and inhibitory connections (Fig. 3a). Inputs projected onto Area 1 units, while outputs were read out from only Area 3 units. We varied the percentage of feedforward connections between areas from 1% to 100%. Networks with sparser feedforward connectivity may result in an information bottleneck that limits communication of color information to downstream areas. Unconstrained networks had color information: even with 1% feedforward connectivity, the color choice could still be decoded from Area 3 units at 81.5% on average, significantly above chance (*p* < 0.01/4). This statistical test is across all trained RNNs, and is only significant when the aggregate mean across all trained networks is reliably above chance. This shows that color representations consistently emerged in unconstrained multi-area networks, so that PMd-like minimal color representations are not solely a consequence of using multiple areas.

**Figure 3:**
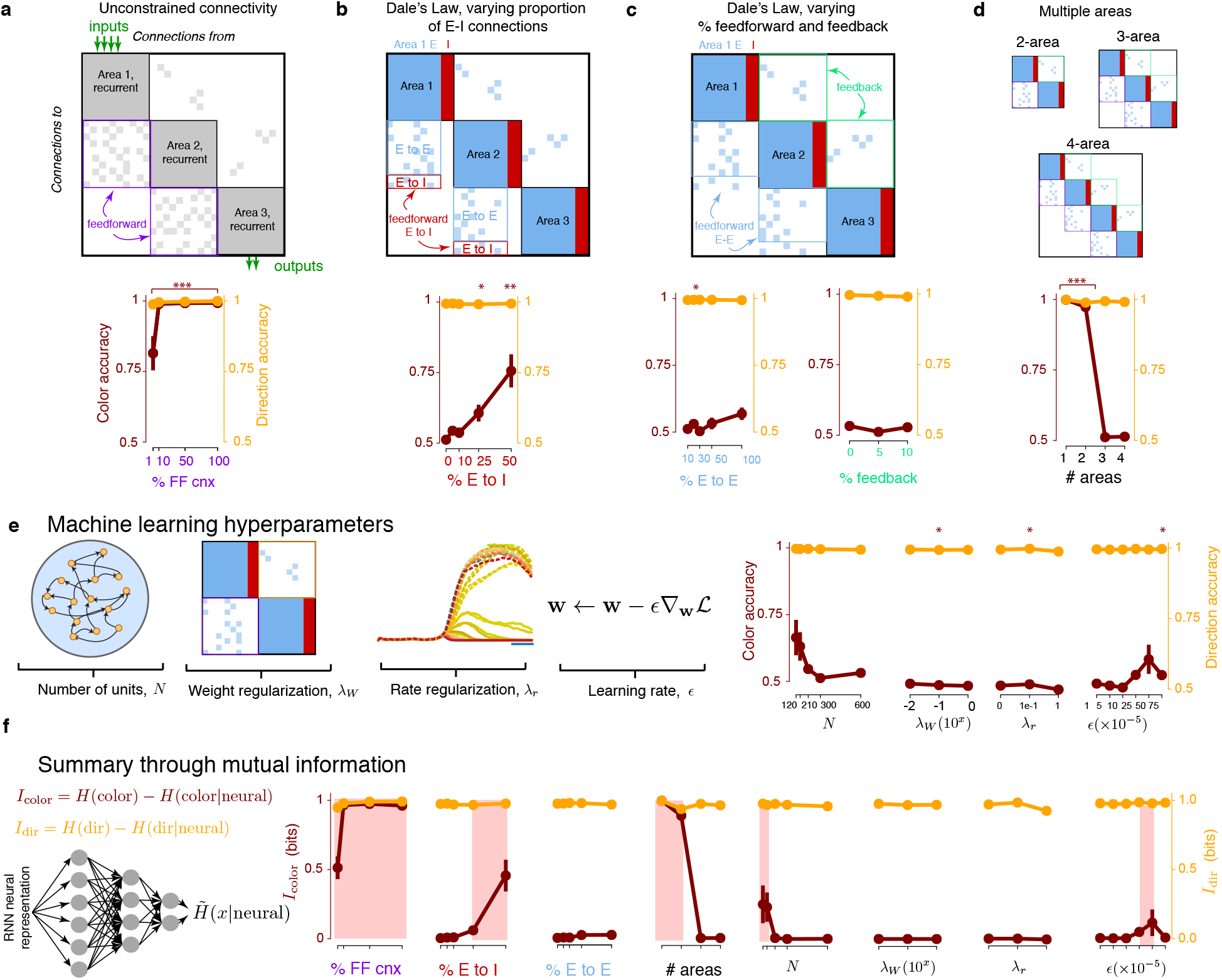
PMd-resembling dynamics emerge in neurologically constrained RNNs. **(a)** We trained 3-area RNNs without explicit excitatory (E) or inhibitory (I) neurons. Inputs projected onto Area 1, and outputs were read out from Area 3. We varied the percentage of feedforward connections and computed the color and direction variance, as well as accuracy, in Area 3. Area 3 had non-zero color variance between 10% to 100% feedforward connections. At 1% feedforward connections, color could still be significantly decoded above chance. Dots are the mean across networks and error bars are s.e.m. For significance, * is *p* < 0.05, ** is *p* < 0.01, and *** is *p* < 0.001 (with appropriate correction for multiple comparisons). **(b)** We incorporated Dale’s law, with 80% E, 20% I neurons, varying the percentage of feedforward E-to-I connections. Minimal representations with chance color decode accuracy emerged when the percentage of feedforward E to I connections (out of all feedforward connections) was 10% or less. **(c)** Color information was relatively robust to feedforward E-E connections. **(d)** At least 3 areas were required for the RNN’s last area to resemble PMd dynamics. **(e)** 3-area RNNs with neurophysiological constraints had minimal representations that were generally robust to machine learning hyperparameters. The only exceptions were when the number of units was relatively small, or the learning rate was relatively large. **(f)** We summarize all sweeps through a mutual information metric, computed by using a variational lower bound on mutual information. We highlighted parameter values in red that lead to stronger color representations in multi-area RNNs.

#### Feedforward E-I connections

We found that neurophysiological architecture constraints were the key design choices leading to RNNs with minimal color representations. The multi-area RNNs presented in Fig. 2 incorporated Dale’s law, with excitatory (80%) or inhibitory (20%) neurons, and excitatory inter-area projections. We identified that the percentage of inter-area feedforward excitatory-to-inhibitory connections (Fig. 3b) strongly impacted minimal color representations. In frontal cortical areas, approximately 10% of units have feedforward connections to downstream areas (Markov et al., 2014), and feedforward excitatory-to-inhibitory connections comprise approximately 10 − 20% of these feedforward connections (Barbas, 2015). In DLPFC in particular, lateral inhibition appears to play a stronger role than feedforward inhibition, contrasting classical sensory circuits where feedforward inhibition is stronger (Datta & Arnsten, 2019). Above 25% feedforward inhibition, color accuracy could be decoded above chance across all trained RNNs (*p* < 0.01/5). At and below 10% feedforward inhibition, color choice decodes achieved near chance accuracies (Fig. 3b, *p* = 0.05, 0.014, and 0.05 for 0%, 5%, and 10% feedforward inhibition). This suggests that low levels of feedforward inhibition result in minimal color representations.

#### E-E connections

Feedforward E-to-E connections had a relatively modest effect on color representation. When there was no feedforward inhibition, we swept the proportion of feedforward E-to-E connections from 10% to 100%, and found that at all levels, there was approximately 0 color variance (Fig. 3c). Across trained RNNs, color could not be decoded significantly above chance levels at all levels (*p* > 0.01/5). We note that at 50% and 100% feedforward E-to-E connections, color representations emerged in a minority of networks, leading to a mean color accuracy above 0.5. However, several networks also had 0.5 decode accuracy. As such, color representations did not reliably emerge at 50% or 100% feedforward E-to-E connectivity. We also varied the percentage of feedback connections at levels of 0%, 5%, and 10%, and found that the last area of these RNNs had minimal color representations.

#### Number of Areas

We observed minimal color representations only when RNNs had 3 or more areas (Fig. 3d,f). 1- and 2-area RNNs trained in the same fashion had above chance color accuracy (Area 1: 1.00, *p* < 0.01/4, Area 2: 0.97, *p* < 0.01/4) in its output area, but 3- and 4- area RNN color decode accuracy was nearly chance (Area 3: 0.51, *p* = 0.05, Area 4: 0.51, *p* = 0.017). This does not mean 1- and 2-area RNNs can never have minimal color representations (see Discussion), but that they do not naturally emerge from optimization with our selected task input and output configuration. We quantified decode accuracy in each sub-area of 4-area RNNs (Fig. S3) and found that color accuracy was near chance levels beginning in Area 3. These results suggest that at least 3-areas are necessary for minimal color representations to naturally emerge in the output area.

#### Machine Learning Hyperparameters

We also tested the effect of machine learning hyperparameters on the emergence of minimal color representations. We varied the number of artificial units, rate regularization, weight regularization, and learning rate (Fig. 3e). We found that RNNs generally had minimal color representations in the last area across hyperparameter sweeps, with two exceptions: when the number of artificial units became too small, or the learning rate became too large, a minority of RNNs had above chance color decode accuracy. This effect was not robust, as several trained RNNs still had nearly chance color decode accuracy (Fig. 3e, *p* > 0.01 for all machine learning hyperparameters). We note that when an RNN has fewer neurons or is trained with a larger learning rate, its learning capacity is reduced (Goodfellow et al., 2016). We summarized all sweeps by computing a variational approximation to the mutual information, which has also been referred to as the usable information (Kleinman et al. (2020), Supplementary Note 2) to quantify the direction and color information in RNNs, as shown in Fig. 3f. This approximation captures how “confident” the decoder was (see Methods), has convenient and interpretable units (bits), and is a variational approximation on mutual information. We observed consistent trends in the effect of hyperparameters when quantifying the mutual information.

#### Training

Finally, we evaluated how color representations change in the network through training. PMd-like 3-area RNNs had nearly zero color information throughout training (Fig. S6a). In contrast, in unconstrained 3-area RNNs, color information increased and plateaued early in training (two representative examples shown in Fig. S6b). While our PMd-like 3-area RNNs outperform the monkeys in accuracy, these results show that had we terminated training earlier, these networks would still have zero color information. In summary, minimal color representations naturally emerged through optimization in multi-area RNNs trained with at least 3 areas, Dale’s law, and connectivity informed by neurobiology.

### Solving the Checkerboard Task: the RNN uses recurrence to compute and output the direction decision on an axis nearly orthogonal to context and color inputs

An RNN is a fully observed system, enabling analysis of its dynamical mechanism. These analyses may produce new mechanistic hypotheses for the computations occurring in areas upstream of PMd, such as the PMdr, DLPFC, and VLPFC. We analyzed how the RNN solves the Checkerboard Task, transforming the targets and checkerboard color input into a direction decision. For simplicity, in the following sections, we refer to the target configuration input as the context input, and the checkerboard input as the color input.

We found that recurrent dynamics play an important role in computation, meaning that the RNN is not strictly input-driven. The Checkerboard Task is analogous to the XOR task, and therefore cannot be solved by a linear transformation of inputs (Supplementary Note 1). We quantified the difference in activity between color decisions (red vs green) and direction decisions (left vs right), calculated as the Euclidean norm between activity for red vs green decisions and left vs right decisions, respectively. This difference was significantly higher in the RNN activity (Fig. 4a, solid lines) than in the RNN input representations (**W**_in_**u**_*t*_, transparent dash-dotted lines) in Areas 1 and 2, suggesting that the RNN amplifies differences in color and direction representations through recurrent dynamics. In Area 3, RNN activity and input representations were similar, suggesting Area 3 is more input-driven than earlier areas, but still uses recurrent dynamics (Area 3 mechanism described later).

**Figure 4:**
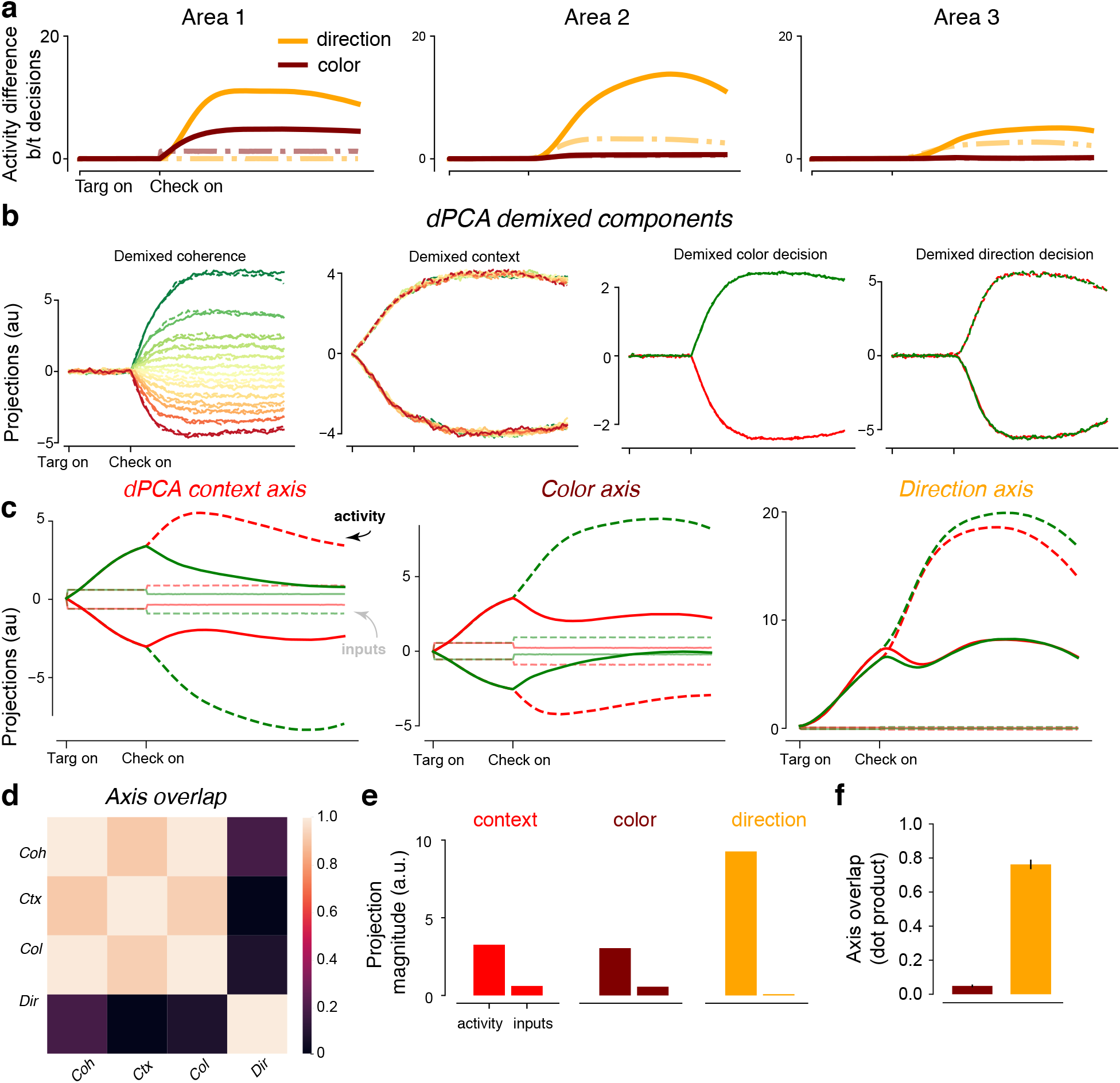
Recurrence and low-dimensional trajectories. **(a)** The activity difference measured as the Euclidean norm between decisions was larger for unit activity (solid) than input representations (transparent dash-dotted) in Areas 1 and 2, suggesting recurrent dynamics play a role in computation these areas. These were comparable in Area 3, suggesting this area is more input driven. **(b)** For Area 1 activity, we performed dPCA and plot the demixed components related to coherence, context (target configuration), color, and direction. **(c)** The context, color, and direction axis correspond to the dPCA principal axes, which are not constrained to be orthogonal. Trajectories for different contexts and colors were separable on both the context and color axis. In contrast, the direction axis separated primarily on chosen direction. The RNN input representation had strong projections on the context and color axes, but not the direction axis. **(d)** The dPCA coherence, context, and color axes were highly overlapping, and orthogonal to the direction axis. **(e)** Summary of the magnitude of activity and input representation projections on each axis. **(f)** Alignment of each axis with the Area 1 to 2 inter-area connections.

How does the RNN represent context and checkerboard inputs, as well as the color and direction choices, to solve the Checkerboard Task? We focused our analysis on Area 1, which uniquely has substantial variance for both color and direction decisions (Fig. 2f), implying a central role in computing the direction choice. We performed dPCA to identify demixed principal components related to the RNN inputs (coherence and context) and decisions (color and direction) (Kobak et al., 2016). We found demixed components that separated information related to coherence, context, the color choice, and the direction choice (Fig. 4b), consistent with these quantities being decodable from activity (Fig. 2g). We subsequently identified the context, color, and direction axes as the dPCA principal axes (unit norm, analogous to PCA eigenvectors), which combine the demixed components (analogous to PCA scores) to reconstruct neural activity (Kobak et al., 2016).

We projected RNN activity and input representations onto the principal axes for context, color, and direction (Fig. 4c). We found that the context and color axis both responded to context and color inputs, and overall trajectories represented both context and color information (Fig. 4c). This suggests that color and context information are mixed in Area 1. In contrast, the direction axis strongly represented the direction choice, but did not strongly represent context or color (Fig. 4c, right). Strikingly, context and color inputs had nearly zero projection on the direction axis (Fig. 4c, right, opaque traces at 0). Consistent with these observations, we found the color and context axes were highly overlapping (dot product: 0.93, Fig. 4d), indicating that context and checkerboard variance are mixed in Area 1 activity. In contrast, the direction axes was closer to orthogonal to the context and color axes (overlap with color and context: 0.14 and 0.09, respectively).

These conclusions were upheld when we performed targeted dimensionality reduction (TDR), where we found (1) a direction axis separating left and right choices, with negligible input projections, and (2) that color and context representations were mixed (Fig. S10). Further, this structure was unique for PMd-like 3-area RNNs. In single-area RNNs, dPCA identified nearly orthogonal context, color, and direction axes, with trajectories that separated almost exclusively based on context, color, and direction, respectively (Fig. S9a).

This Area 1 representation has an important property: the direction choice is represented robustly on a nearly orthogonal axis that has close to zero context and color input projections, (Fig. 4e). This is not trivial: as counter-examples, single-area RNNs use orthogonal direction axes that have context and color input projections (Fig. S9a), while an unconstrained 3-area RNN direction axis has context and color information, and also receives context and color inputs (Fig. S9b).

We hypothesized this representation is beneficial for minimal color representations: to propagate direction, but not color information, the inter-area connections from Area 1 to 2 would align with the direction axis, which does not represent color information or inputs. We therefore expect the Area 1 to 2 readout to be highly aligned with the direction axis, but not aligned with the context or color axis. We found the direction axis was preferentially aligned with the Area 1 readout (mean overlap: 0.76 ± 0.03. (mean ± s.e.m. Fig. 4f), while the color axis was not (overlap: 0.05 ± 0.01.). Together, these results show that Area 1 computes and outputs the direction choice through recurrent dynamics on an axis that: (1) has negligible input projections, and (2) is highly aligned with the Area 1 readout.

### Solving the Checkerboard Task: a dynamical mechanism

What dynamical mechanism does the RNN employ to compute and output a direction choice that is orthogonal to RNN inputs? Since the inputs have nearly zero projection on the direction axis, there is no direct integration of the inputs on the direction axis in Area 1. We visualized the dynamics of the RNN (Kao, 2019) in the PCs to determine the mechanism used by Area 1 to solve the Checkerboard Task. In all analyses in this section, we show the PCs and dynamics of the pre-activation RNN state, denoted by **x** (and dynamics 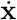), described in the Methods.

We analyzed RNN dynamics in the top 2 PCs, which capture 97.7% of the variance. As shown in Fig. 5a, in the Targets On epoch, the RNN state diverges to two attractor regions with slow dynamics, corresponding to the two target configurations. A checkerboard input delivered in this attractor region causes the RNN state to diverge to 4 trajectories, corresponding to each possible combination of color and direction choice. We projected the dPCA color, context, and direction principal axes onto these PCs, shown in Fig. 5b. We also projected the two possible context inputs (pink) and checkerboard inputs (red or green) on the PCs, which drive RNN activity along the color and context axis (Fig. 5c). Irrespective of the target configuration, green checkerboards cause the RNN state to increase along PC_2_ while red checkerboards cause the RNN state to decrease along PC_2_. The strength of the input representation is state-dependent: checkerboards corresponding to left reaches, whether they are green or red, cause smaller movements of the RNN state along the color axis.

**Figure 5:**
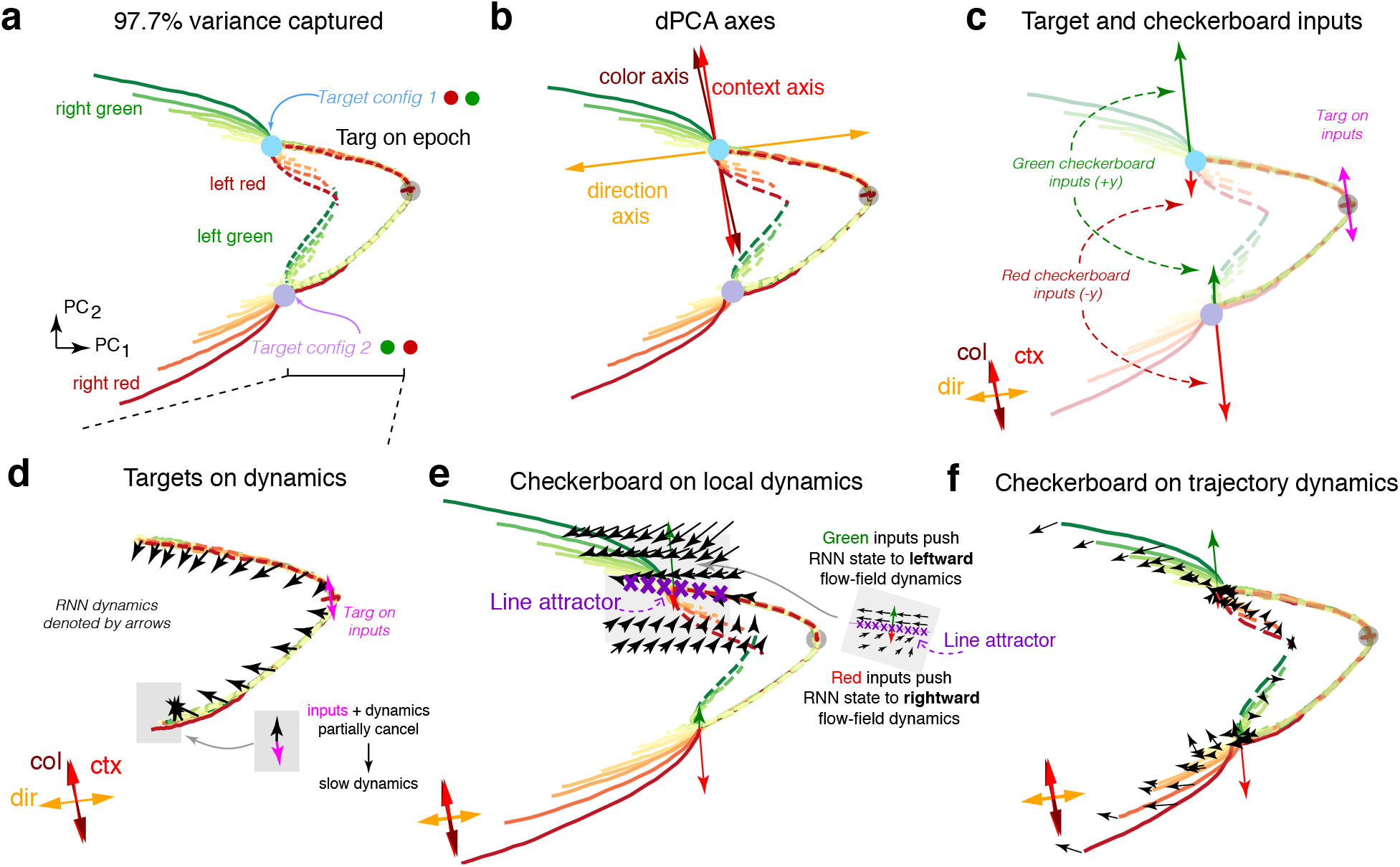
A dynamical mechanism for computing the direction choice. **(a)** Top 2 PCs of Area 1 activity, which capture 97.7% of the Area 1 variance. In the targets on epoch, the trajectories separate to two regions corresponding to the two potential target configurations (Target config 1 in blue, and Target config 2 in purple). The trajectories separate upon checkerboard color input, leading to four total trajectory motifs: right green, left red, right red, and left green. **(b)** Projection of the dPCA principal axes onto the PCs. **(c)** Projection of the context and color inputs onto the PCs. Context inputs are shown in pink, a green checkerboard input in green, and a red checkerboard input in red. Green (red) checkerboards lead to an increase (decrease) in PC_2_ and the color axis, and differ in magnitude depending on the location of the trajectory in PC space. Trajectories are reduced in opacity to better visualize inputs. **(d)** Visualization of RNN dynamics and inputs. We visualize RNN dynamics by projecting them into the PCs, visualized as a flow field (black). These dynamics and inputs drive the trajectories to one of two attractor regions with slow dynamics, where the inputs and dynamics destructively interfere. **(e)** At the target config 1 attractor, a green (red) checkerboard increases (decreases) the RNN state along PC_2_ and color axis. We plotted the local dynamics around this region using our previously described technique (Kao, 2019). We found that upwards green inputs pushed the RNN trajectories to a leftward flow field (increasing activity along direction axis), while red inputs pushed the RNN trajectories into a rightward flow field (decreasing activity along direction axis). These recurrent dynamics enable activity to change significantly on the direction axis, while having nearly zero input projections. These flow fields were separated by a line attractor. Arrows are not to scale; checkerboard inputs have been amplified to be visible. **(f)** We visualized the dynamics across multiple trajectory motifs. Arrows are not to scale, for visualization purposes.

We visualized the dynamics, 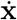, by projecting them into the PCs. In the Targets On epoch, shown in Fig. 5d, context inputs cause movement along the context axis. The RNN dynamics implemented a leftward flow-field that pushed the RNN state into an attractor region of slow dynamics. At the end of the Targets On epoch, the RNN dynamics point in a direction largely opposing the context inputs, destructively interfering and resulting in slow dynamics (Fig. 5d, inset). The combination of context inputs and RNN dynamics result in activity along the context, color, and direction axes (compare to Fig. 4d).

We next visualized RNN dynamics when the Checkerboard input was delivered, leading to a direction choice (Kao, 2019). We found that RNN dynamics were oriented to either increase or decrease along the direction axis depending on the RNN’s state. As shown in Fig. 5e, focusing on the Target config 1 attractor, green checkerboards cause upwards movement, while red checkerboards cause downwards movement. The RNN implements approximately opposing flow fields above and below the attractor. Above the attractor, a leftward flow-field increases direction axis activity, while below the attractor, a rightward flow-field decreases direction axis activity. A green checkerboard input therefore pushes the RNN state into the leftward flow-field (solid green trajectories) while a red checkerboard input pushes the RNN state into a rightward flow-field (dotted red trajectories). This computes the direction choice in a given context, while allowing the direction axis to be orthogonal to color inputs. These dynamics hold in both target configurations, as shown in Fig. 5f, leading to separation of right and left decisions on the direction axis.

Further analysis of the network’s dynamics revealed a line attractor (Fig. 5e), likely reflecting an evidence integration process (Mante et al., 2013). The line attractor identified was oriented between the color inputs and the direction axis in the PCs (Fig. S7). Above the line attractor, recurrent dynamics drive the PC trajectory to the left, while below the line attractor, dynamics drive the trajectory to the right (Fig. 5e). If inputs are turned off, the trajectory relaxes to the line attractor, reflecting integrated evidence (Mante et al., 2013). These results add to a growing literature, where integration along a line attractor is a common computational motif observed in RNNs that integrate evidence towards binary decisions (Mante et al., 2013, Maheswaranathan et al., 2019, Maheswaranathan & Sussillo, 2020).

### Inter-area connections propagate direction and filter color information

Our analyses in a previous section (Fig. 4f) suggest that the direction axis is preferentially aligned with the readout axis whereas color and context are not. We more fully characterized the inter-area connections between RNN areas. We denote the feedforward connections from Area 1 to 2 as **W**_12_, and from Area 2 to 3 as **W**_23_. We present results for feedforward connections from excitatory connections from excitatory units. Based on the hypothesis that the brain uses null and potent spaces to selectively filter information (Kaufman et al., 2014), we evaluated the potent and null spaces of **W**_12_ and **W**_23_. We defined the potent space to be the right singular vectors corresponding to the largest singular values (see Methods). The null space corresponded to the singular vectors with the smallest singular values.

We quantified how the color and direction axis were aligned with these potent and null spaces (see Methods). The projections onto the potent space are shown in Fig. 6a,b for **W**_12_ and **W**_23_, respectively. The null projection magnitudes are equal to one minus the potent projection. We found the direction axis was more aligned with the potent space and the color axis was more aligned with the null space. In fact, the direction axis was most aligned with the top singular vector, whereas the color axis was similarly aligned to a random vector. This alignment was robust to the dimension of the effective potent space, and was consistent across networks with varying feedforward connectivity percentages (10%, 20%, 30%, 50%, 100%). This indicates that direction information is preferentially propagated to subsequent areas, while color information is not. This phenomena is schematized in Fig. 6c.

**Figure 6:**
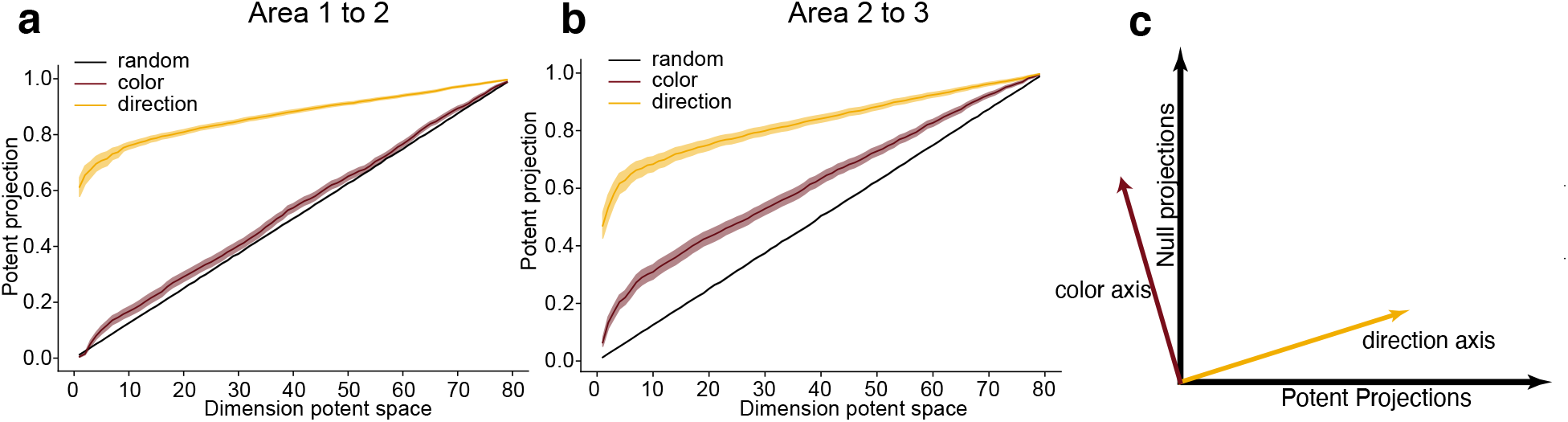
**(a)** Projections onto the potent space between Areas 1 and 2 for the color and direction axis, and a random vector as a function of effective rank for the input area to the middle area. Regardless of the dimension of the potent space, the direction axis is preferentially aligned with the potent space, indicating the information along this axis propagates, while the color axis is more aligned with the nullspace. Shading indicates s.e.m. **(b)** Same as (a) but for projections between Areas 2 and 3. **(c)** Illustration depicting how the orientation of the axes affect the information that propagates.

These results also have implications on how inter-area connections relay information between areas. Color activity has significant representation in Area 1 (see Fig. 2). Therefore, the inter-area connections must not merely propagate the highest variance dimensions of a preceding area (Semedo et al., 2019). Consistent with this hypothesis, we found that while the top 2 PCs capture 97.7% excitatory unit variance, the top 2 readout dimensions of **W**_12_ only captured 40.0% of Area 1’s excitatory unit neural variance (Fig. S12). Hence, inter-area connections are not aligned with the most variable dimensions, but are rather aligned to preferentially propagate certain types of information — a result consistent with a recent study analyzing links between activity in V1 and V2 (Semedo et al., 2019).

We also quantified the alignment of feedback connections with the color and direction axes in each Area, using networks with 5% feedback connectivity. We found that between Areas 2 and 1, both the direction and color axes had significantly above-chance alignments with feedback connections, with the direction axis having the highest alignment (Fig. S13). Between Areas 3 and 2, the direction and color axes had significant above chance alignment, but the effect was relatively small. Recall that the color axis in Area 2 and 3 captures almost zero variance. These results indicate that Area 1, which computes the direction decision, also receives direction inputs from downstream areas.

### Area 3, modeling PMd dynamics, is primarily input driven and implements bistable dynamics

We showed previously that Area 3 most closely resembled PMd’s dynamics (Fig. 2). Our results suggest that a direction signal has been computed before Area 3 and is selectively propagated through the RNN’s inter-area connections. We found that the input to Area 3 (through **W**_23_) is a graded direction signal that provides a directional evidence signal for left or right reaches (Fig. 7a). This activity must be transformed into eventual DV outputs, which are the accumulated evidence for a left or right reach. This is illustrated in Fig. 7a, where we plot 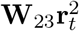 (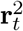 are the unit activations of Area 2), and 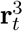.

**Figure 7:**
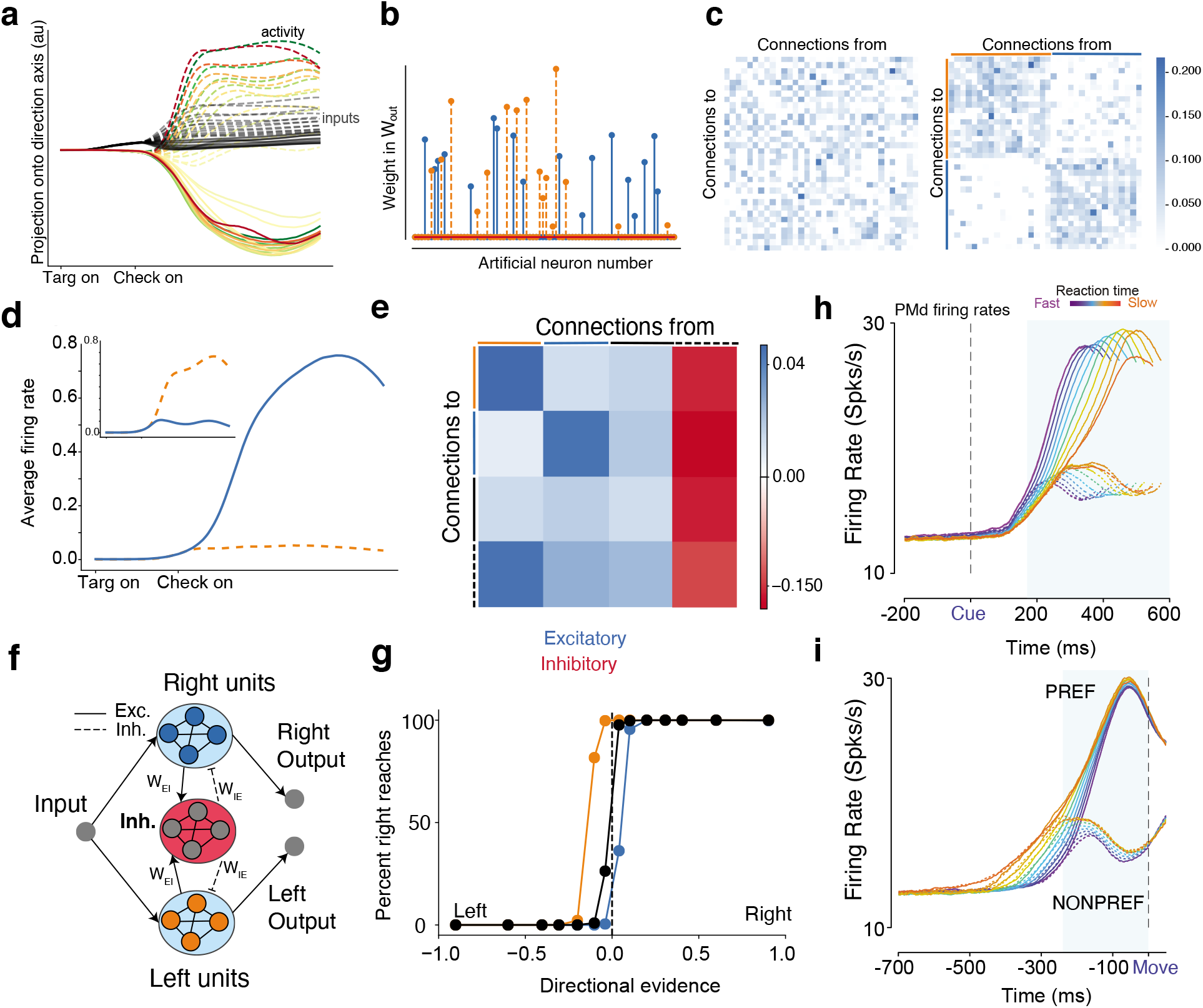
Area 3 mechanism. **(a)** Projection of input and overall activity onto the direction axis identified through dPCA. The conventions are the same as in Figure 4. **(b)** Readout weights in **W**_out_ are sparse, with many zero entries, and selective weights for a left or right reach. **(c)** The unsorted connectivity matrix for the nonzero readout units (left panel), and the sorted connectivity matrix when the matrix was reordered based on the readout weight pools (right). **(d)** Average PSTHs from units for a leftward reach and (inset) rightwards reach. When one pool increases activity, the other pool decreases activity. **(e)** Averaged recurrent connectivity matrix. **(f)** Schematic of output area. **(g)** Psychometric curve after perturbation experiment, where 10% of inhibitory weights to the left pool (orange) and right pool (blue) were increased (doubled). Directional evidence is computed by using the signed coherence and using target configuration to identify the strength of evidence for a left reach and strength of evidence for a right reach. Increasing inhibition to the left excitatory pool leads to more right choices and vice versa. **(h,i)** Firing rates of PMd neurons for PREF direction reaches and NONPREF direction reaches aligned to checkerboard and movement onset. In Winner-take-all models, when one pool wins the firing rate of the other pool is suppressed due to lateral inhibition. In PMd, when the PREF direction wins the firing rate of the NONPREF direction decreases (blue shaded region in i).

To analyze Area 3’s dynamics, we first observed that **W**_out_’s coefficients were sparse, with 44 out of 80 output weights being identically zero. We found that the readout led to two separate clusters of artificial units: units with non-zero coefficients for the left DV (orange) and those with non-zero coefficients for the right DV (blue). Artificial units projected either to the left or right DV outputs, but not both, suggesting that there are two clusters mediating left and right choices.

Based on this clustering, we sorted and visualized the excitatory units of **W**_rec_, which upon first glance generally has no discernible structure (Fig. 7c, left panel). After sorting, we found that two self-excitatory pools of units emerged in **W**_rec_, the first pool in Fig. 7c (right) corresponding to the left DV and the second pool corresponding to the right DV. In addition to these two pools, we identified a pool of randomly connected excitatory units and a pool of inhibitory units with strong projections from and to the two pools. The full connectivity matrix is shown in Fig. S14. This structure is consistent with a winner-take-all network, where increasing activity in one pool inhibits activity in the other pool through a separate inhibition pool (Fig. 7d). By taking the averaged connectivity matrix, similar to Yang et al. (2019), we confirmed that there were two excitatory pools that received similar projections from the random excitatory pool and inhibitory pool (Fig. 7e). We summarize the behavior with a schematic of the area in Fig. 7f.

We subsequently applied selective perturbations to **W**_rec_ to determine how behavioral performance was biased. We increased inhibition to either the right or the left pool by doubling the weights of 10% of the inhibitory neurons associated with each pool. We found that this biased the network towards more left or right reaches, respectively, shown in Fig. 7g. When inhibition was increased to the right excitatory pool, the network was more likely to respond left. Conversely, when inhibition was increased to the left excitatory pool, the network was more likely to respond right.

When analyzing PMd neurons, sorted based on their preferred and non-preferred directions, we found that preferred reaches amplified activity while non-preferred reaches suppressed activity, consistent with winner-take-all dynamics (see dip in NONPREF firing rates aligned to checkerboard, Fig. 7h, and before movement initiation in Fig. 7i). Activity here is sorted by RT and choice, and similar effects were seen when sorting by other variables such as coherence. Together, these results show that the output area, modeling PMd, robustly transforms separable direction inputs to a decision variable through a winner-take-all mechanism.

## Discussion

Even though behavior and cognition arise from the coordinated computations of multiple brain areas, there is limited understanding of how interacting brain areas coordinate to produce cognitive behavior (Semedo et al., 2019, Kohn et al., 2020). In this study, we used multi-area RNNs to gain mechanistic insight into how the brain computes a perceptual decision in the Checkerboard Task and transmits only the direction decision to PMd. These results propose hypotheses for computations that occur upstream of PMd, particularly how neural population activity representing context, color, and direction are structured, and what information is propagated between areas.

Our analysis provided two key insights for how multi-area RNNs compute based on coordinated within-area dynamics and inter-area connections. First, we observed that Area 1 used a dynamical mechanism to separate the direction information on a direction axis. Color and context inputs had a negligible projection onto this direction axis, but the combination of inputs and recurrent dynamics led to separation of activity along the direction axis (Fig. 5e,f). This critical feature, that color inputs and direction outputs were decoupled through recurrent dynamics, facilitated propagation of direction – and not color – information to downstream areas. Second, inter-area connections were preferentially aligned to the direction axis, not axes of maximal variance, leading to selective propagation of direction axis activity and attenuation of color axis activity. This role for inter-area connections is consistent with null and potent spaces for filtering and propagating information between areas (Kaufman et al., 2014, Stavisky et al., 2017) and communication subspaces, which are aligned with lower variance dimensions (Semedo et al., 2019).

Our task could be solved with or without feedback connections with equivalent performance, indicating that feedback was not necessary to solve the task (Fig. S8). When the model had feedback connections, we observed that feedback connections between Areas 2 and 1 preferentially conveyed direction information. One interpretation is that these feedback inputs play a role in computing the eventual direction decision. Another perspective on feedback signals is that they may related to error signals used for learning (Lillicrap et al., 2020). Multi-area networks may help understand and develop new hypotheses for physiological studies of feedforward and feedback computation (Cumming & Nienborg, 2016, Nienborg & Cumming, 2009), and more generally distributed processing for decision-making and cognition (Pinto et al., 2019, Kauvar et al., 2020). Future research may use carefully designed tasks in conjunction with multi-area RNNs to better understand the role of feedback in computation.

Our results suggest that cortex and multi-area RNNs may share a more general principle of multi-area information processing: if information becomes irrelevant for later computations, it is reduced. In the Checkerboard Task, color information is necessary to compute the direction decision, but does not need to be represented after the direction decision is computed, as in PMd (Chandrasekaran et al., 2017, Wang et al., 2019, Yamagata et al., 2012, Cisek & Kalaska, 2005, Coallier et al., 2015). In deep neural networks, it is believed that minimal representations simplify the role of the output classifier (Achille & Soatto, 2018, Soatto & Chiuso, 2016). This idea is consistent with (1) the multi-area RNN developing a minimal (little color information) but sufficient (robust direction information) representation of task inputs, and (2) Area 3, the output area, using a simple winner-take-all readout, forming two pools of neurons representing right and left decisions (Fig. 7).

We found there were important design considerations in training multi-area RNNs to resemble PMd activity. PMd-like minimal color representations naturally and robustly emerged through optimization in RNNs trained with at least 3 areas, Dale’s law, and neurophysiological architectural constraints. 4-area networks also had no color information by Area 3 (Fig. S3). It is worth noting that RNN training time generally increased with more areas. Future work should assess how learning with these architectural constraints and backpropagation-through-time lead to PMd-like activity and dynamics. Our initial experiments suggest that neurophysiological constraints strongly bias the networks to learn solutions that only contain direction information, and further that color information was not present in the network during training (Fig. S6).

Our analysis of RNNs trained with different architectural and neurophysiological constraints also suggest that the strength of feedforward E-I and E-E connections can have a profound role in shaping the computations in these RNNs. Based on these results, we speculate that in the brain, altering the relative strength of feedforward E-I connections might allow cortical networks to control information flow and computations (Large et al., 2016). For instance, in sensory areas, increased feedforward connections may allow the rapid transmission of information from one area to the next. In contrast, to simplify downstream readouts and also to increase recurrent processing within areas, modulating relative levels of recurrent excitation/inhibition, feedforward, and feedback connections may allow for better control of information flow (Datta & Arnsten, 2019). We believe this is a rich avenue for future work in both experimental and theoretical contexts.

We chose 3 areas in our multi-area RNN, which was the minimal number of areas needed for PMd-like representations to naturally emerge. We emphasize that we are not suggesting that 3 areas are ideal or necessary for solving the task; indeed, 4-area RNNs also have minimal color representations. The pathway likely involved in solving this task includes the VLPFC, DLPFC, PMdr, and PMdc (Hoshi, 2013). The VLPFC is directedly connected to the inferotemporal cortex through the uncinate fasciculus (Yeterian et al., 2012, Gerbella et al., 2009) and sends projections to PMd via DLPFC and PMdr (Takahara et al., 2012). Our 3-area RNN should therefore be viewed as a demonstration of the importance of incorporating multiple areas in an RNN and as a model that contains the key features of multi-area computation. Our analyses investigating the alignment of information with inter-area connections can, in principle, be extended to an arbitrary number of areas. Ultimately, the number of areas used for modelling neurophysiological tasks should depend on experiments (Yamagata et al., 2012, Siegel et al., 2015, Pinto et al., 2019) and anatomical studies (Barbas & Pandya, 1987, Luppino et al., 2003, Takahara et al., 2012).

Single-area RNNs trained with the same task configuration and hyperparameters had significant color variance and information across many hyperparameter settings (Fig. S5). In no way should these results be construed as a rejection of the usefulness of a single-area RNN model for modeling cognition. Single-area RNNs can still model PMd, for example, by changing the task inputs to be signed directional evidence instead of checkerboard color (Mante et al., 2013). As a proof of concept, we trained a single-area RNN to process signed directional evidence, which by definition did not have color information (Fig. S15). This RNN, consistent with prior work, integrated inputs along the direction readout, and had winner-take-all structure, like Area 3 (Williams et al., 2018).

In general, RNN computations can be strongly affected by the input (Remington et al., 2018, Kao, 2019) and output configuration. Although using signed directional evidence inputs into a single-area RNN provides some insight into PMd dynamics, it uses human expertise to design inputs that remove color information. Such an RNN trained with these inputs cannot propose a mechanism for how context and color evidence are transformed into a direction decision, or how these representations propagate through inter-area connections. In contrast, because PMd-like dynamics emerge through multi-area RNN optimization, we were able to discover a new mechanism for computing the direction decision and study how coordinated dynamics and inter-area connections propagate direction information and attenuate color information. Thus, single-area RNN models will provide critical insight when they are used to study local RNN computations (Mante et al., 2013, Sussillo et al., 2015, Michaels et al., 2016, Song et al., 2016, Chaisangmongkon et al., 2017, Remington et al., 2018). In contrast, multi-area RNNs can complement such insights and provide holistic insight into distributed computations. We believe this is increasingly important, as neurophysiologists are recording from multiple brain areas (Steinmetz et al., 2018, 2019, Pinto et al., 2019).

While they bear resemblance, Area 1 of our multi-area RNN differs from the RNN trained in Mante et al. (2013). We note that while the task the animals solved in Mante et al. (2013) was an extension of the Checkerboard task, their network and neural data analyses combined color and motion evidence into a signed directional evidence signal. Thus, in Mante et al. (2013)’s model, they solved a multiplexing operation, where depending on the context, the network selected and integrated one input (signed-directional color or motion evidence) but not the other (Table S2). In contrast, our network solved the color-to-direction transformation, which is the XOR problem, where depending on the context (target configuration), the color input (checkerboard) resulted in a different direction output (Table S1). This led to a related, but different computation. In Mante et al. (2013), the network solved the multiplexing operation with dynamics such that a selection vector selected the relevant input and ignored the other. Our network did not use a selection vector. Instead, all inputs were relevant and needed to be nonlinearly combined to compute the direction output. Depending on target configuration, the same color inputs moved the trajectory into flow fields leading to different movement along the direction axis, thereby solving the XOR task. Further, the Area 1 network solved this task in a way such that downstream areas had a minimal representation of color information, resembling PMd. This required the Area 1 computation to (1) use precisely oriented inputs and dynamics in Area 1 (Fig. 5e) that nearly orthogonalized color inputs and direction output, and (2) preferentially propagate the direction output to downstream areas through alignment with inter-area connections. These aspects of the mechanism are unique to multi-area RNNs, reflecting a distributed computation, in contrast to Mante et al. (2013).

An important advantage of RNN modeling is that probing an RNN enables mechanism discovery and produces testable hypotheses for future experiments. By analyzing the RNN’s activations and parameters, we were able to discover a mechanism for computing a direction decision in the Checkerboard Task, investigate the role of inter-area connections, and propose how an input-driven output area produces PMd resembling dynamics. Our analysis of the multi-area RNN leads to testable hypotheses for future experiments. First, cortical areas upstream of PMd should exhibit mixed selectivity for color and direction information, consistent with studies of DLPFC in cognitive tasks (Rigotti et al., 2013, Mante et al., 2013, Fusi et al., 2016). Neural population dynamics should diverge to two regions with slow dynamics based on target configuration, with largely overlapping context and color axes, but an orthogonal direction axis. Second, due to alignment of inter-area connections, direction axis activity in DLPFC should be more predictive of activity in downstream regions such as PMdr and PMd than activity in the top PCs.

## Methods

### Somatomotor reaction time visual discrimination task and recordings from PMd

The task, training and electrophysiological methods used to collect the data used here have been described previously Chandrasekaran et al. (2017) and are reviewed briefly below. All surgical and animal care procedures were performed in accordance with National Institutes of Health guidelines and were approved by the Stanford University Institutional Animal Care and Use Committee.

#### Task

Two trained monkeys (Ti and Ol) performed a visual reaction time discrimination task Chandrasekaran et al. (2017). The monkeys were trained to discriminate the dominant color in a central static checkerboard composed of red and green squares and report their decision with an arm movement. If the monkey correctly reached to and touched the target that matched the dominant color in the checkerboard, they were rewarded with a drop of juice. This task is a reaction time task, so that monkeys initiated their action as soon as they felt they had sufficient evidence to make a decision. On a trial-by-trial basis, we varied the signed color coherence of the checkerboard, defined as (*R* − *G*)/(*R* + *G*), where R is the number of red squares and G the number of green squares. The color coherence value for each trial was chosen uniformly at random from 14 different values arranged symmetrically from 90% red to 90% green. Reach targets were located to the left and right of the checkerboard. The target configuration (left red, right green; or left green, right red) was randomly selected on each trial. Both monkeys demonstrated qualitatively similar psychometric and reaction-time behavior. Results in Fig. 1b,c correspond to Monkey Ti.

For analyses in Fig. 7g, we combined signed coherence with the target configuration to estimate a parameter we term signed directional evidence. For instance, a completely red checkerboard with a signed coherence of 1 and a red target on left (green target on right) would be considered to have −1 directional evidence. Conversely, if the target configuration was such that the red target was on the right (green target on left), the directional evidence would be 1.

#### Recordings

In our original study, we reported the activity of 996 units recorded from Ti (n=546), and Ol (n=450) while they performed the task (Chandrasekaran et al., 2017). PMd units demonstrate temporal complexity in their firing rate profiles. In Chandrasekaran et al. (2017), we applied the visuomotor index of that measures the degree of sustained activity to separate this population into the broad categories of increased, perimovement and decreased units. Increased units exhibited ramp-like increases in average firing rate, time-locked to the onset of the visual stimulus, with slopes that varied with stimulus coherence. These are the classic candidates for neurons that might carry an integrated decision variable for the discrimination task. Monkey Ol and Ti’s PMd units both had low choice color probability. Reported analyses from PMd data use units pooled across Monkey Ol and Ti.

### RNN description and training

We trained a continuous-time RNN to perform the checkerboard task. The RNN is composed of *N* artificial neurons (or units) that receive input from *N*_in_ time-varying inputs **u**(*t*) and produce *N*_out_ time-varying outputs **z**(*t*). The RNN defines a network state, denoted by 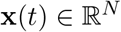; the *i*th element of **x**(*t*) is a scalar describing the “currents” of the *i*th artificial neuron. The network state is transformed into the artificial neuron firing rates (or network rates) through the transformation:

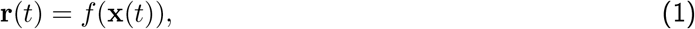

where *f*(·) is an activation function applied elementwise to **x**(*t*). The activation function is typically nonlinear, endowing the RNN with nonlinear dynamics and expressive modeling capacity Goodfellow et al. (2016). In this work, we use *f*(*x*) = max(*x*,0), also known as the rectified linear unit, i.e., *f* (*x*) = relu(*x*). In the absence of noise, the continuous time RNN is described by the equation

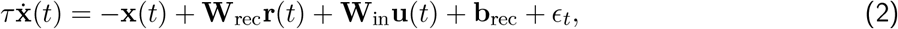

where *τ* is a time-constant of the network, 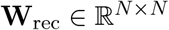 defines how the artificial neurons are recurrently connected, 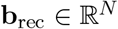 defines a constant bias, 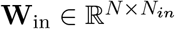 maps the RNN’s inputs onto each artificial neuron, and *ϵ_t_* is the recurrent noise. The output of the network is given by a linear readout of the network rates, i.e.,

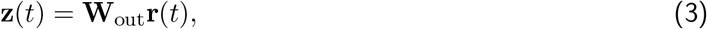

where 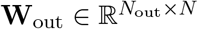 maps the network rates onto the network outputs.

We trained RNNs to perform the checkerboard task as follows. For all networks, unless we explicitly varied the amount of units, we used *N*_in_ = 4, *N* = 300, and *N*_out_ = 2.

The four inputs were defined as:

1. Whether the left target is red (−1) or green (+1).
2. Whether the right target is red (−1) or green (+1).
3. Signed coherence of red (ranging from −1 to 1), (*R* − *G*)/(*R* + *G*).
4. Signed coherence of green (ranging from −1 to 1), (*G* − *R*)/(*R* + *G*). Note that, prior to the addition of noise, the sum of the signed coherence of red and green is zero.

The inputs, 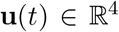, were defined at each time step, *t*, in distinct epochs. In the ‘Center Hold’ epoch, which lasted for a time drawn from distribution 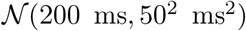, all inputs were set to zero. Subsequently, during the ‘Targets’ epoch, which lasted for a time drawn from distribution 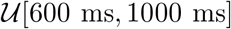, the colors of the left and right target were input to the network. These inputs were noiseless, as illustrated in Fig. 2a, to reflect that target information is typically unambiguous in our experiment. Following the ‘Targets’ epoch, the signed red and green coherences were input into the network during the ‘Decision’ epoch. This epoch lasted for 1500 ms. We added zero mean independent Gaussian noise to these inputs, with standard deviation equal to 5% of the range of the input, i.e., the noise was drawn from 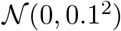. At every time point, we drew independent noise samples and added the noise to the signed red and green coherence inputs. We added recurrent noise 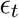, adding noise to each recurrent unit at every time point, from a distribution 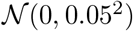. Following the ‘Decision’ epoch, there was a ‘Stimulus Off’ epoch, where the inputs were all turned to 0.

The two outputs, 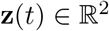 were defined as:

1. Decision variable for a left reach.
2. Decision variable for a right reach.

We defined a desired output, **z**_des_(*t*), which was 0 in the ‘Center Hold’ and ‘Targets’ epochs. During the ‘Decision’ epoch, **z**_des_(*t*) = 1. In the ‘Stimulus Off’ epoch, **z**_des_(*t*) = 0. In RNN training, we penalized output reconstruction using a mean-squared error loss,

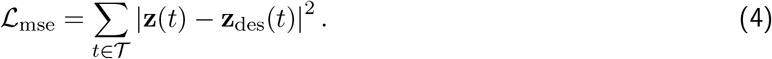

The set 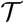 included all times from all epochs except for the first 200 ms of the ‘Decision’ epoch from the loss. We excluded this time to avoid penalizing the output for not immediately changing its value (i.e., stepping from 0 to 1) in the ‘Decision’ epoch. Decision variables are believed to reflect a gradual process consistent with non-instantaneous integration of evidence, e.g., as in drift-diffusion style models, rather than one that steps immediately to a given output.

To train the RNN, we minimized the loss function:

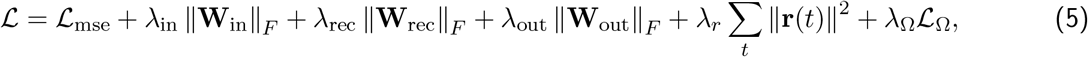

where

▪ ∥**A**∥_*F*_ denotes the Frobenius norm of matrix **A**
▪ λ_in_ = λ_rec_ = λ_out_ = λ*_r_* = 1 to penalize larger weights as well as rates (Sussillo et al., 2015, Michaels et al., 2016).
▪ λ_Ω_ = 2
▪ 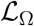 is a regularization term that ameliorates vanishing gradients proposed and is described in prior literature (Pascanu et al., 2013, Song et al., 2016).

During the training process, we also incorporated gradient clipping to prevent exploding gradients (Pascanu et al., 2013). Training was performed using stochastic gradient descent, with gradients calculated using backpropagation through time. For gradient descent, we used the Adam optimizer, which is a first order optimizer incorporating adaptive gradients and momentum (Kingma & Ba, 2014).

We stopped training to prevent the network from achieving nearly perfect performance on the task. Prior studies modeling cognitive tasks have employed early stopping to reproduce animal behavior Mante et al. (2013), Song et al. (2016). Every 200 or 500 training epochs, we generated 2800 cross-validation trials, 100 for each of the 28 possible conditions (14 coherences × 2 target configurations). For each trial, there was a correct response (left or right) based on the target configuration and checkerboard coherence. When training, we defined a “correct decision” to be when the RNNs DV for the correct response was greater than the other DV and the larger DV was greater than a pre-set threshold of 0.6. We evaluated the network 500ms before the checkerboard was turned off (the end of the trial). We required this criteria to be satisfied for at least 65% of both leftward and rightward trials. We note that this only affected how we terminated training. It had no effect on the backpropagated gradients, which depended on the mean-squared-error loss function.

When testing, we defined the RNNs decision to be either: (1) whichever DV output (for left or right) first crossed a pre-set threshold of 0.6, or (2) if no DV output crossed the pre-set threshold of 0.6 by the end of the ‘Decision epoch,’ then the decision was for whichever DV had a higher value at the end of this epoch — an approach that is well established in models of decision-making (Brunton et al., 2013, Ratcliff, 1978). If the RNN’s decision on a single trial was the same as the correct response, we labeled this trial ‘correct.’ Otherwise, it was incorrect. The proportion of decisions determined under criterion (2) was negligible (0.5% across 100 trials for each of 28 conditions using the RNN shown in Fig. 2). An interpretation for criterion (2) is that if the RNN’s DV has not achieved the threshold certainty level by the end of a trial, we assign the RNN’s decision to be the direction for which its DV had the largest value. Finally, in training only, we introduced ‘catch’ trials 10% of the time. On 50% of catch trials, no inputs were shown to the RNN and **z**_des_(*t*) = 0 for all *t*. On the remaining 50% of catch trials, the targets were shown to the RNN, but no coherence information was shown; likewise, **z**_des_(*t*) = 0 for all *t* on these catch trials.

We trained the three-area RNNs by constraining the recurrent weight matrix **W**_rec_ to have connections between the first and second areas and the second and third areas. In a multi-area network with N neurons and m areas, each area had N/m neurons. In our 3-area networks, each area had 100 units. Of these 100 units, 80 were excitatory and 20 were inhibitory. Excitatory units were constrained to have only positive outgoing weights, while inhibitory units were constrained to have only negative outgoing weights. We used the pycog repository Song et al. (2016) to implement these architecture constraints. The parameters for the exemplar RNN used in the paper are shown in Table 1. In our hyperparameter sweeps, we varied the hyperparameters of the exemplar RNN. For each parameter configuration, we trained 8 different networks with different random number generator seeds.

**Table 1:** Hyperparameters of exemplar RNN.

| Hyperparameter | Value |
| --- | --- |
| Number of units | 300 |
| Number of areas | 3 |
| Learning rate | 5e-5 |
| Time Constant | 50ms |
| Discretization bin width | 10ms |
| Rate regularization | 0 |
| Weight regularization | 1 |
| Activation function | Relu |
| Feedforward connection | 10% |
| Feedback connections | 5% |
| Dale law | Yes |

### Decoding analysis for PMd data

For PMd data, we calculated decoding accuracy using 400 ms bins. We report numbers in a window [−300ms, +100 ms] aligned to movement onset. We used the MATLAB *classify* command with 75% training and 25 % test sets. Decoding analyses were performed using 5-31 simultaneously recorded units from Plexon U-probes and the averages reported are across 51 sessions. To assess whether decoding accuracies were significant on a session by session basis, we shuffled the labels 200 times and estimated the 1^st^ and 99^th^ percentiles for this surrogate distribution. The decode accuracy for direction, color, and context variables for a session was judged to be significant if it lay outside this shuffled accuracy. Every session had significant direction decode, while no session had significant color and context decode accuracy.

### Decoding and Mutual information for RNNs

We used a decoder and mutual information approximation to quantify the amount of information (color, context, direction) present in the network. We trained a neural network to predict a relevant choice (for example, color) on a test set from the activity of a population of units. We used 700 trials for training, and 2100 independent trials for testing. To generate the trials for training and testing, we increased the recurrent noise to be drawn from the distribution 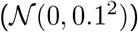 to prevent overfitting. For each trial, we averaged data in a window [−300ms, +100ms] around reaction time.

We trained a neural network with 3 layers, 64 units per layer, leakyRelu activation (*α*=0.2), and dropout (p=0.5), using SGD, to predict the choice given the activity of the population. We removed the leakyRelu activation for the linear network, and increased dropout (p=0.8). For both the nonlinear and linear network, we trained the neural network to minimize the cross-entropy loss. We used the same neural network from the decode to compute an approximation to mutual information, described in Supplementary Note 2.

### RNN behavior

To evaluate the RNN’s psychometric curve and reaction-time behavior, we generated 200 trials for each of the 28 conditions, producing 400 trials for each signed coherence. For these trials, we calculated the proportion of red decisions by the RNN. This corresponds to all trials where the DV output for the red target first crossed the preset threshold of 0.6; or, if no DV output crossed the threshold of 0.6, if the DV corresponding to the red target exceeded that corresponding to the green target. The reaction time was defined to be the time between checkerboard onset to the first time a DV output exceeded the preset threshold of 0.6. If the DV output never exceeded a threshold of 0.6, in the reported results, we did not calculate a RT for this trial.

### Visualizing RNN dynamics

To visualize RNN dynamics in Fig. 5d,f, we computed the RNN’s state dynamics, 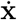, along selected PSTHs, and subsequently projected onto the PCs. To visualize the local RNN dynamics in Fig. 5e, we used the technique described in Kao (2019), where we sampled dynamics by modifying the RNN state along the axes defined by PCs 1 and 2. As described further in our prior study (Kao, 2019), there are important considerations in visualizing dynamics in PCs, since trajectory movement in the orthogonal dimension can dramatically change the dynamics. As such, Fig. 5e should be viewed as a figure for intuition, while Fig. 5f definitively show the sampled dynamics in support of the proposed mechanism.

### dPCA

Demixed principal components analysis (dPCA) is a dimensionality reduction technique that provides a projection of the data onto task related dimensions while preserving overall variance (Kobak et al., 2016). dPCA achieves these aims by minimizing a loss function:

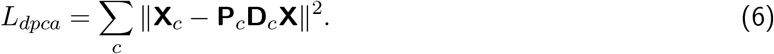

Here, ***X**_c_* refers to data averaged over a “dPCA condition” (such as time, coherence, context, color, or direction), having the same shape as 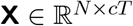, but with the entries replaced with the condition-averaged response. The aim is to recover (per dPCA condition *c*) a **P**_*c*_ and **D**_*c*_ matrix. **P**_*c*_ is constrained to have orthonormal columns, while **D**_*c*_ is unconstrained. The number of columns of **P**_*c*_ and rows of **D**_*c*_ reflects the number of components one seeks to find per condition. We project the data onto the principal components **D**_*c*_**X** to observe the demixed components (Fig. 4b). The column of **P**_*c*_ reflects how much the demixed data contributes to each neuron. We use the principal axes from **P**_*c*_ to compute the axis overlap, as in Kobak et al (Kobak et al., 2016). We used axes of dimension 1 for RNNs, which were sufficient to capture most color, context, or direction variance. For the neural data Fig. 1e, we used five components for direction, color and context since the PMd data was higher dimensional than the RNNs.

Our results were consistent if we used dPCA or TDR (Fig. S10). The top principal axis from each **P**_*c*_ are analogous to the axes found from TDR. Both methods seek to reconstruct neural activity from demixed components. To apply TDR, one explicitly parametrizes task variables (See Targeted Dimensionality Reduction in Methods), while **D**_*c*_**X** serves the purpose of finding demixed components in dPCA. Overall, the choice of using dPCA or TDR to find the axes did not affect our conclusions.

For multi-area analyses, we separated the units for each area and found the task-relevant axes for this subset of units. For the inter-area analyses, we used RNNs with only excitatory connections, and therefore found the color and direction axis using only the excitatory units (Fig. S11). In all other analyses, all units were used to identify the axes. For RNN activity, we performed dPCA using activity over the entire trial. For PMd activity, we used a window of (0ms, 800ms) relative to checkerboard onset. We restricted time windows for the PMd activity because we wanted to minimize movement related variance.

### Targeted Dimensionality Reduction

Targeted dimensionality reduction (TDR) is a dimensionality reduction technique that finds low dimensional projections that have meaningful task interpretations. We applied TDR as described by the study by Mante, Sussillo and colleagues (Mante et al., 2013). We first z-scored the firing rates of each of the 300 units across time and trials, so that the firing rates had zero mean and unit standard deviation. We then expressed this z-scored firing rate as a function of task parameters using linear regression,

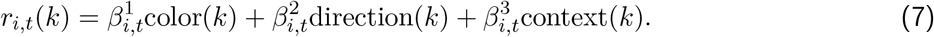

Here, *r_i,t_*(*k*) refers to the firing rate of unit *i* at time *t* on trial *k*. The total number of trials is *N*_trials_. This regression identifies coefficients 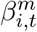 that multiply the m^th^ task parameter to explain *r_i,t_*(*k*). We defined the task parameters as follows:

▪ color(*k*) was the signed coherence of the checkerboard on trial *k*, given by (*R* − *G*)/(*R* + *G*).
▪ direction(*k*) was −1 for a left decision and +1 for a right decision.
▪ context(*k*) was the target orientation, taking on −1 if the green (red) target was on the left (right) and +1 if the green (red) target was on the right (left).

We did not fit a bias term since the rates were z-scored and therefore zero mean. For each unit, *i*, we formed a matrix **F**_*i*_ having dimensions *N*_trials_ × 3, where each row consisted of [color(*k*), direction(*k*), context(*k*)]. We define **r***_i,t_* to be the rate of unit *i* at time *t* across all trials. We then solved for the coefficients, denoted by 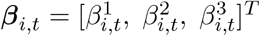, using least squares,

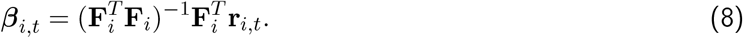

Each ***β**_i,t_* is therefore a 3 × 1 vector, and concatenating ***β**,_i,t_* across *t* results in ***β**_i_*, a 3 × *T* matrix, of which there are *N*. We then formed a tensor where each *β_i_* is stacked, leading to a tensor with dimensions 3 × *T* × *N*. For each of the 3 task variables, we found the time *T* where the norm of the regression coefficients, across all units, was largest. For the *m*th task variable, we denote the vector 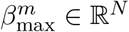 to be a vector of coefficients that define a 1-dimensional projection of the neural population activity related to the *m*th task variable. These vectors are what we refer to as the task related axes. To orthogonalize these vectors, we performed QR decomposition on the stacked ***β***_max_ matrix 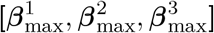, which is an *N* × 3 matrix. This decomposition finds orthogonal axes so that the axes would capture independent variance.

### Choice probability

To calculate the choice probability for a single unit, we first computed the average firing rate in a window from [−300 ms, +100 ms] around the reaction time for each trial. We used the average firing rates calculated across many trials to create a firing rate distribution based on either the color decision (trials corresponding to a red or green choice) or the direction decision (trials corresponding to a left or right choice).

To compute the color choice probability, we constructed the firing rate distributions corresponding to a green choice or red choice. If these two distributions are non-overlapping, then the neuron has a color choice probability of 1; the average firing rate will either overlap with the red or green firing rate distributions, but not both. On the other hand, if the two distributions are completely overlapping, then the neuron has a color choice probability of 0.5; knowing the firing rate of the neuron provides no information on whether it arose from the red or green firing rate distribution. When there is partial overlap between these two distributions, then firing rates where the distributions overlap are ambiguous. We computed choice probability as the area under the probability density function at locations when the two distributions did not overlap, divided by 2 (to normalize the probability). To calculate the direction choice probability, we repeated the same calculation using firing rate distributions corresponding to a left choice or right choice.

### Canonical correlation

We applied CCA to assess the similarity between neural activity and the artificial unit activity (Sussillo et al., 2015). Before applying CCA, we performed principal component analysis to reduce the dimensionality of the artificial and neural activity to remove noise (Sussillo et al., 2015). We reduced the dimensionality to 3 and 8 for RNNs and PMd, respectively. These dimensionalities were chosen as they captured over 88% of the variance for each dataset when aligned to checkerboard. We report the average CCA correlation coefficients in Fig. 2 using times in a window of [0, 400ms] aligned to checkerboard onset for the PMd and RNN activity. The data was binned in 10ms bins.

### Axis overlap

To calculate axis overlap (Fig. 4h), we first note that the direction, color, and readout axes were unit length vectors. When vectors are unit length, their dot product is equal to the cosine of the angle between them, which is 1 for parallel vectors and 0 for orthogonal vectors. We therefore quantified axis overlap by taking the dot product between the direction (color) axis and the readout axis. We note that the readout axis only received projections from excitatory units, so we performed dPCA to find the color and direction axis using only the excitatory units.

### Analyses of inputs and activity

In order to disentangle the effects of external inputs and recurrence, in Fig. 4a, we evaluated the input contribution and overall activity. For Area 1, we defined the input contribution as **W**_in_**u**_*t*_, and for areas 2 and 3, we defined the input contribution as 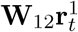, and 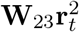 respectively, where 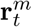 denotes the activity of the units in area *m*. The activity 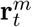 corresponds to the firing rate that experimentalists could measure, reflecting a combination of input and recurrent interactions. For constant inputs, a stable value of the activity implies there is little recurrent processing.

### Null and Potent Analyses

To calculate the overlap between the color and direction axes with the null and potent spaces, we performed singular value decomposition on the inter-area connections, **W**_12_ and **W**_23_. **W**_12_ and **W**_23_ were 80 × 80 matrices, and were full rank. Nevertheless, they had near some zero singular values, indicating that the effective rank of the matrix was less than 80. We defined the potent dimensions to be the top *m* right singular vectors, while the null dimensions were the remaining 80 − *m* right singular vectors.

We performed the analyses of Fig. 6a,b by varying the potent and null dimensions, sweeping *m* from 1 to 80. For each defined potent and null space, we calculated the axis overlap between the direction (or color) axis and the potent (or null) space by computing the L2-norm of the orthogonal projection (squared). We report the squared quantity because the expectation of the norm of a projection of a random vector onto an m-dimensional subspace of an n dimensional space is *m/n*. We include an approximation of the expectation of the projection of a random vector in Fig. 6a,b by averaging the projection of 100 random vectors. Our results show that the direction axis was always more aligned with potent dimensions than the color axis, irrespective of the choice of *m*, and that the direction axis was preferentially aligned with the top singular vector.

### Visualization of neural activity in a low dimensional space

The activity of multiple units on a single trial is high dimensional, with dimension equal to the number of units. To visualize the activity in a lower dimensional space, dimensionality reduction techniques can be used. In addition to TDR, we also utilized Principal Components Analysis (PCA) and t-distributed Stochastic Neighbor Embedding (tSNE) to visualize neural activity in low-dimensional spaces.

PCA finds a linear low-dimensional projection of the high dimensional data that maximizes captured variance. We performed PCA on both the experimental data and RNN rates. PCA is an eigenvalue decomposition on the data covariance matrix. To calculate the covariance matrix of the data, we averaged responses across conditions. This reduces single trial variance and emphasizes variance across conditions. Firing rates were conditioned on reach direction and signed coherence. In both the experimental data and RNN rates, we had 28 conditions (14 signed coherences each for left and right reaches).

tSNE embeds high dimensional data in a low dimensional manifold that is nonlinear, enabling visualization of activity on a nonlinear manifold. The tSNE embedding maintains relative distances between data points when reducing dimensionality, meaning that points closer in high dimensional space remain closer when viewed in a low dimensional manifold. We projected our data into a two dimensional manifold. We used the default parameters from the sickit-learn implementation (https://scikit-learn.org/stable/modules/generated/sklearn.manifold.TSNE.html). The data visualized under tSNE was averaged in a window [−300ms, +100ms] around reaction time.

## Supplementary Figures, and Notes

**Figure S1:**
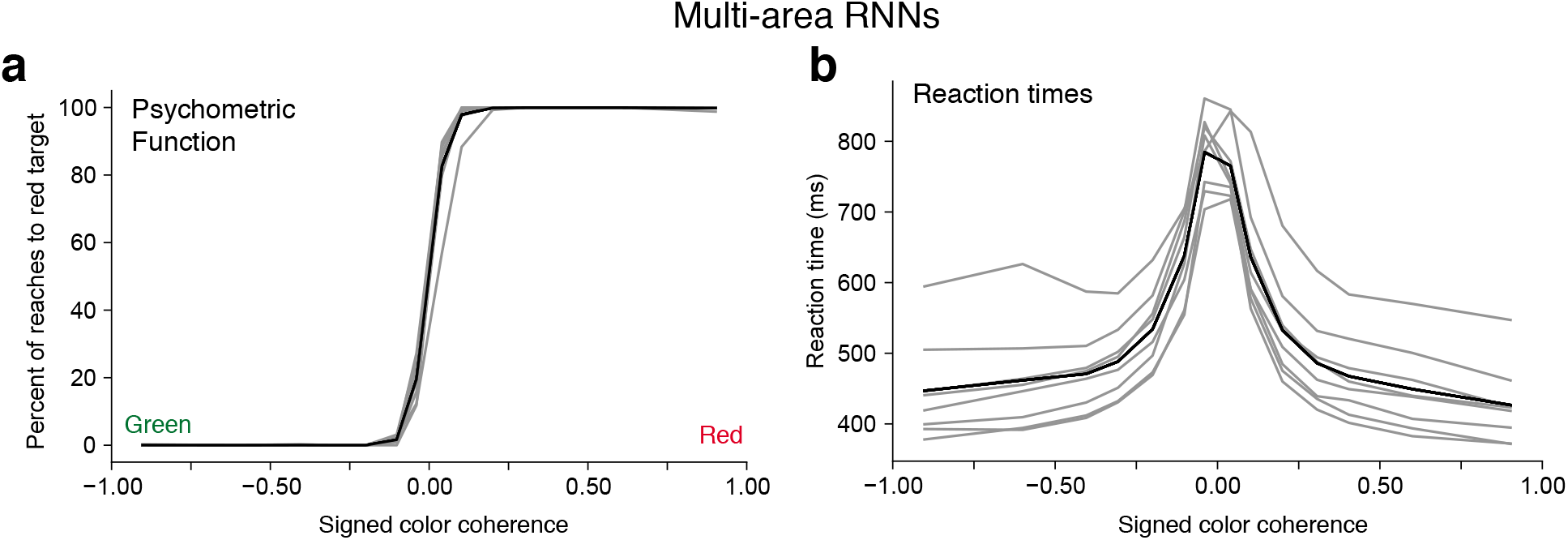
(**a**) Psychometric and (**b**) reaction time curves across multi-area (right) RNNs with Dale’s law trained for this study. The hyperparameters used for these RNNs are described in Table 1. Gray lines represent individual RNNs and the black solid line is the average across all RNNs.

**Figure S2:**
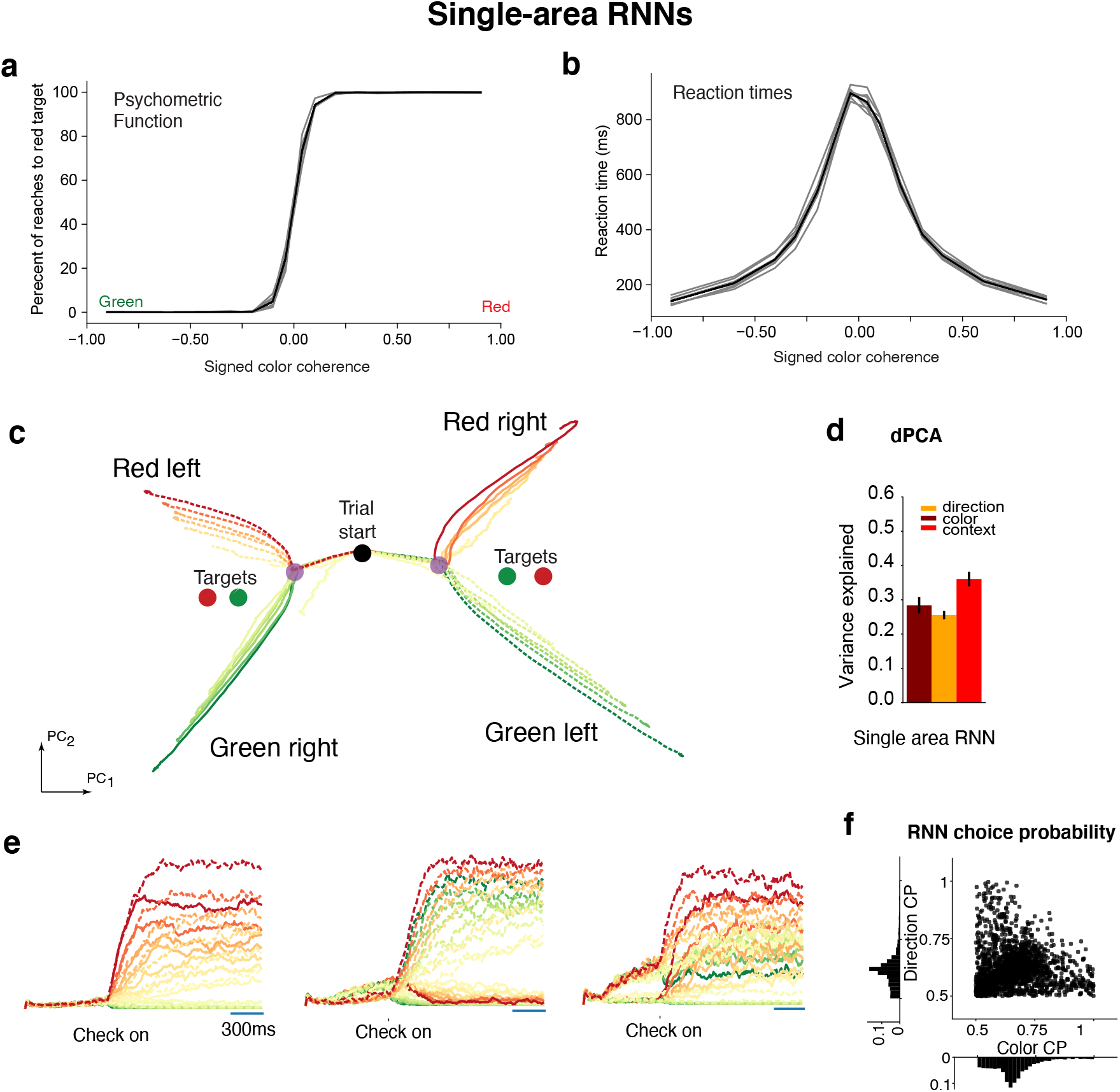
Single-area RNN dynamics. **(a)** Psychometric curves for single-area RNNs trained with Dale’s Law to solve the checkerboard task. The inputs and outputs of these networks were the same as the multi-area RNNs in Fig. 2. **(b)** Chronometric curves for the single-area RNNs shown in a. **(c)** Single-area RNN neural trajectories in the top 2 PCs. Single-area RNNs had four trajectory motifs for each combination of (left vs right) and (red vs green). In the Targets epoch, the RNN’s activity approached one of two locations in state space (purple dots), corresponding to the two target configurations. In the checkerboard epoch, trajectories separate based on the coherence of the checkerboard, causing 4 total distinct trajectory motifs. Although the direction decision is not separable in the principal components, the direction decision is separable in higher dimensions (see the direction axis found using dPCA in Fig. S9a). **(d)** dPCA variance captured for the color (28%), context (26%), and direction (36%) axes for the RNN. **(e)** Example RNN PSTHs. (Left) Primarily coherence selective, (middle) primarily direction selective, and (right) mixed selectivity. **(f)** Choice probability for simulated single-area RNN units. Unlike PMd choice probabilities (Fig. 1g), many units have high color choice probabilities.

**Figure S3:**
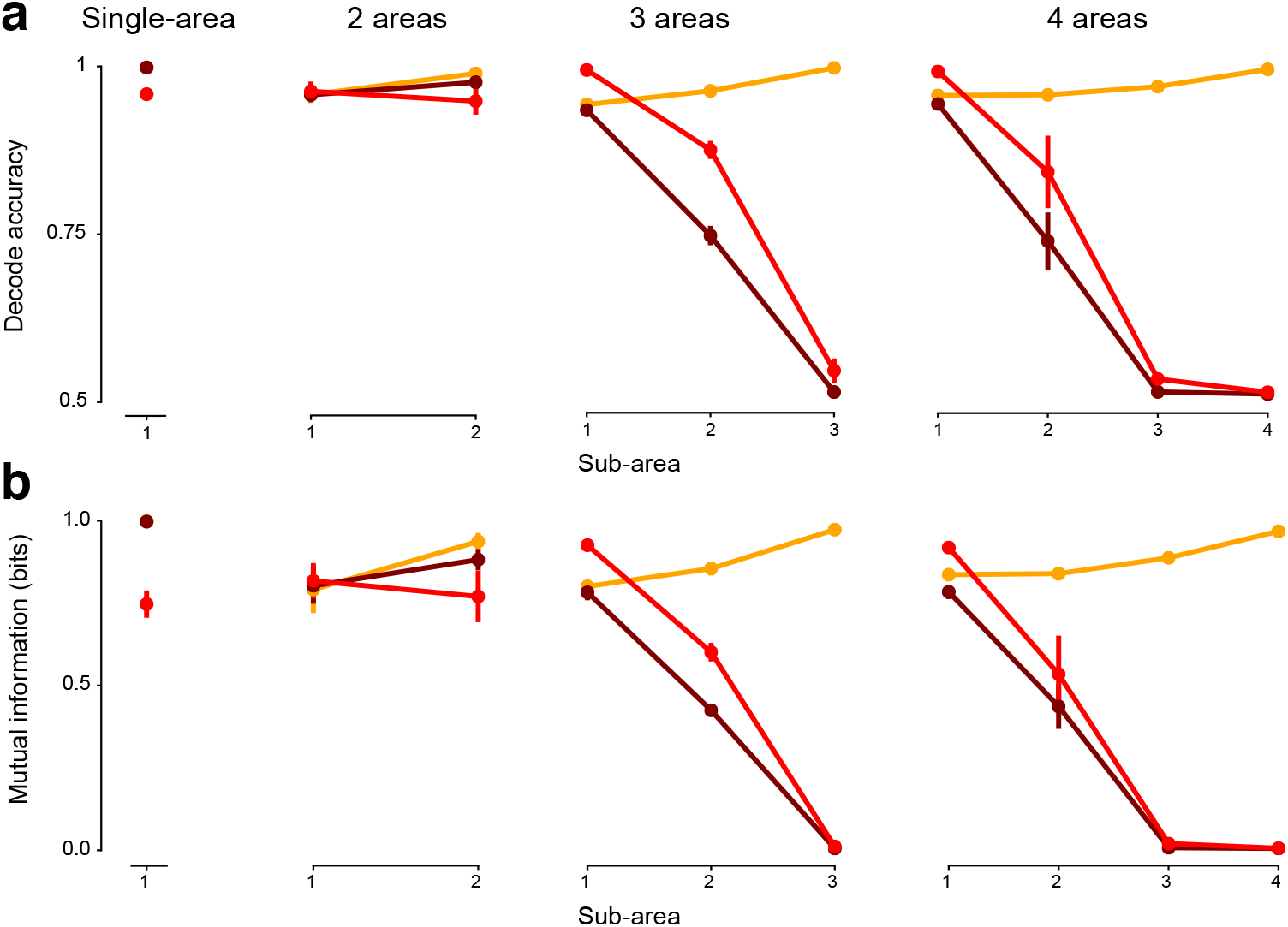
Decode accuracy and mutual information per area in multi-area RNNs. **(a)** Decode accuracy in each area for 1- to 4-area RNNs for color, context, and direction corresponding to Fig 3d. The 3- and 4-area RNNs had minimal color representations in their last area. Note that the 4-area RNN also has a minimal color representation in Area 3. **(b)** Mutual information in each area for 1- to 4-area RNNs. Color conventions as in Fig. 3. Red is context, dark brown is color, and orange is direction.

**Figure S4:**
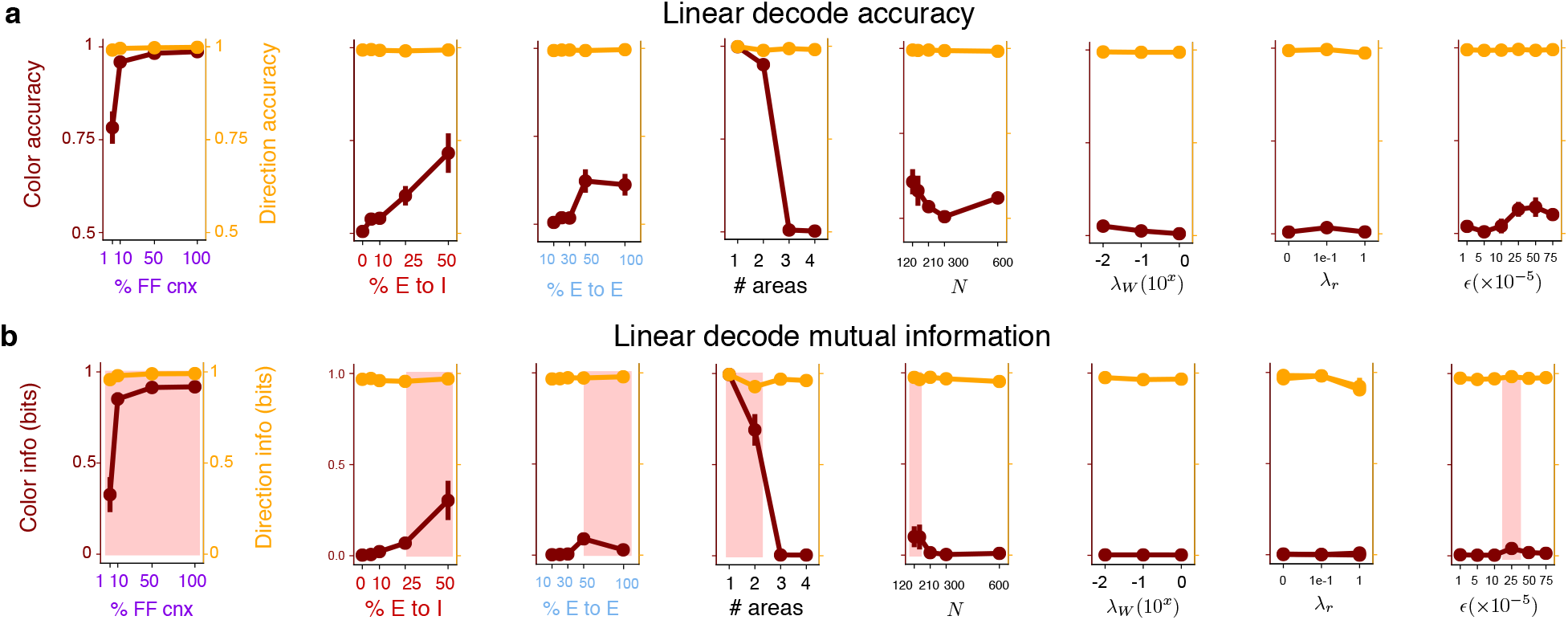
Results of Fig. 3 reproduced with a linear classifier. This figure reproduces the simulations in Fig 3, but with a linear classifier. The main conclusions are upheld. **(a)** Linear decode accuracy for all hyperparameter sweeps shown in Fig. 3. **(a)** Mutual information estimated by using the linear network trained with cross entropy loss.

**Figure S5:**
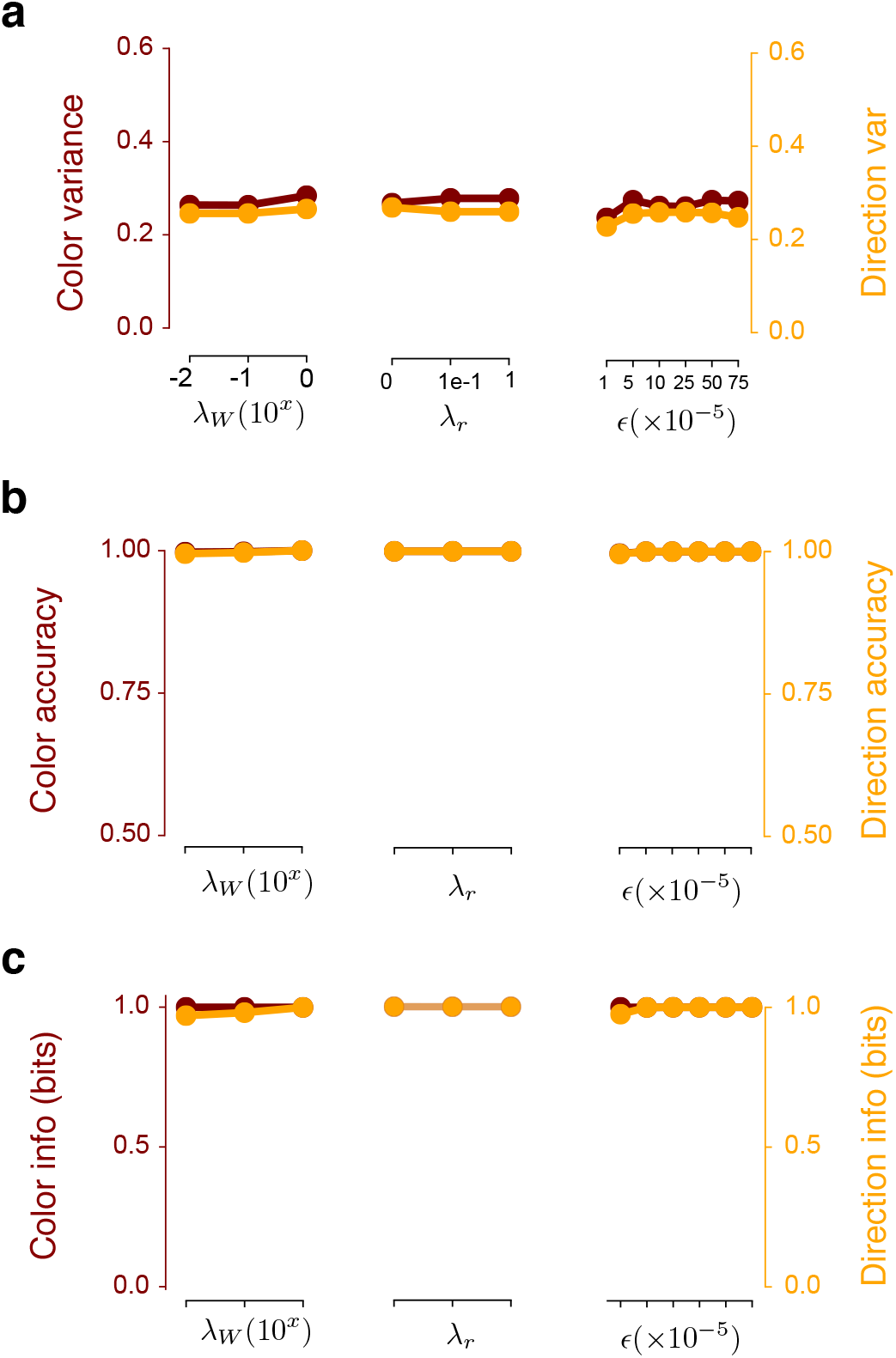
Hyperparameter sweeps for single-area RNNs. **(a)** dPCA color and direction variance captured for three different regularization parameters (weight regularization: λ*_w_*, rate regularization: λ*_r_*, and learning rate: *ϵ*). There is a significant color representation in all optimized single-area RNNs. **(b)** Decode accuracy of the color and direction decision; color accuracy is at 1 (hidden behind direction accuracy) for the three different hyper parameters. The color decode accuracy (maroon) is at nearly 1 across all tested hyperparameters. These points are behind the direction decode accuracy (orange). **(c)** Mutual information estimate. The color mutual information (maroon) is nearly 1 across all tested hyperparameters.

**Figure S6:**
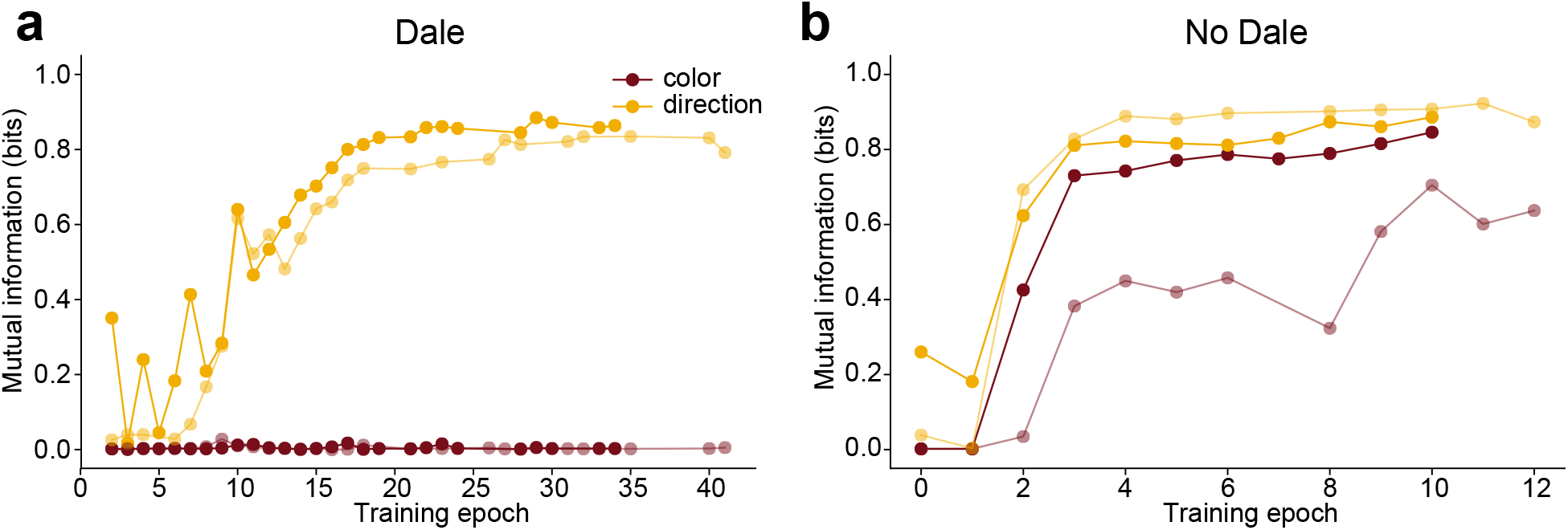
Color and direction information through training in Area 3. Each “training epoch” represents 500 iterations of gradient descent. **(a)** In the PMd-like 3-area RNNs that were trained with Dale’s law, color information in Area 3 remained near zero throughout training (two different representative networks, light and dark shade). **(b)** In the unconstrained 3-area RNNs, color information in Area 3 increased early in training and appeared to plateau (two different networks, light and dark shade). Networks were only saved if the loss function decreased, so certain training epochs are not present.

**Figure S7:**
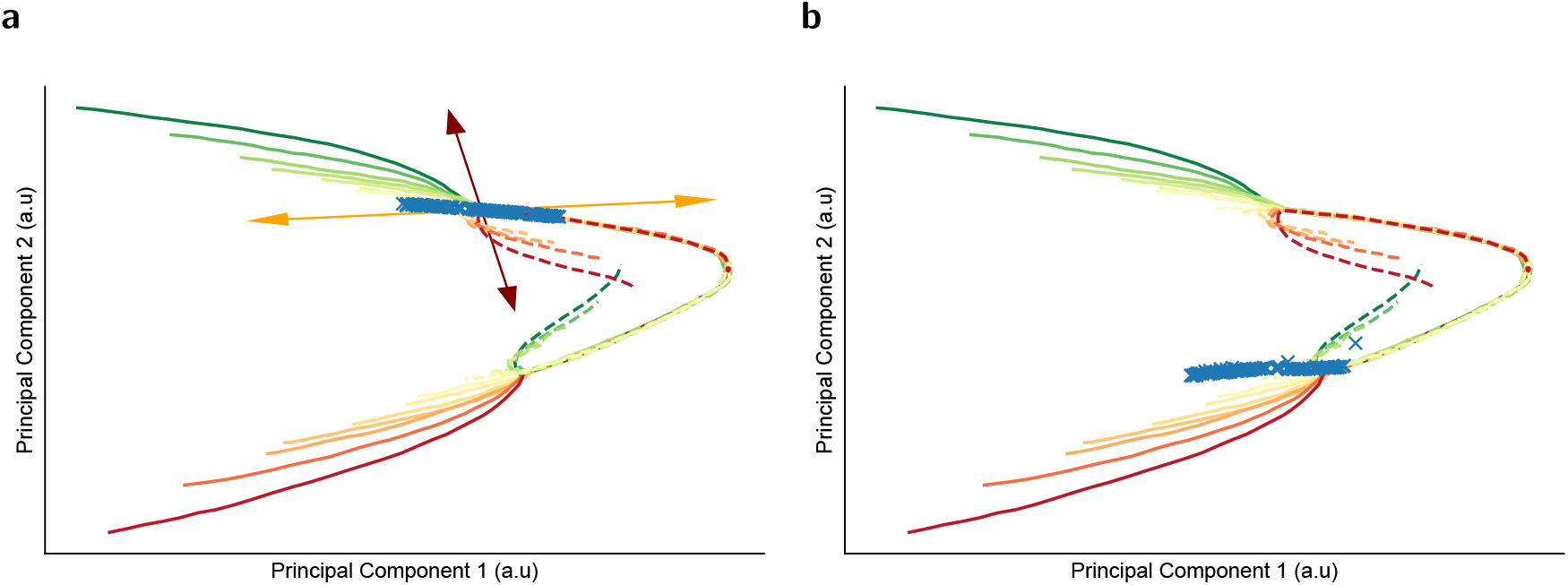
**(a)** We identified slow points (blue crosses, which are areas where the RNN dynamics 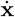 satisfy 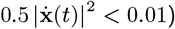) for Area 1 found in one context (green right, red left). The slow points are shown overlapped with the principal component activity. Slow points were found with the code associated with a recent review paper (Yang & Wang, 2020). When finding the slow points, the context inputs were left on and the checkerboard inputs were turned off (equivalent to Mante et al. (2013), where the context input was on but motion and color evidence were off). In the exemplar network, we found a line attractor during the decision epoch, consistent with prior literature (Mante et al., 2013). **(b)** Slow points identified for the other context form another approximate line attractor.

**Figure S8:**
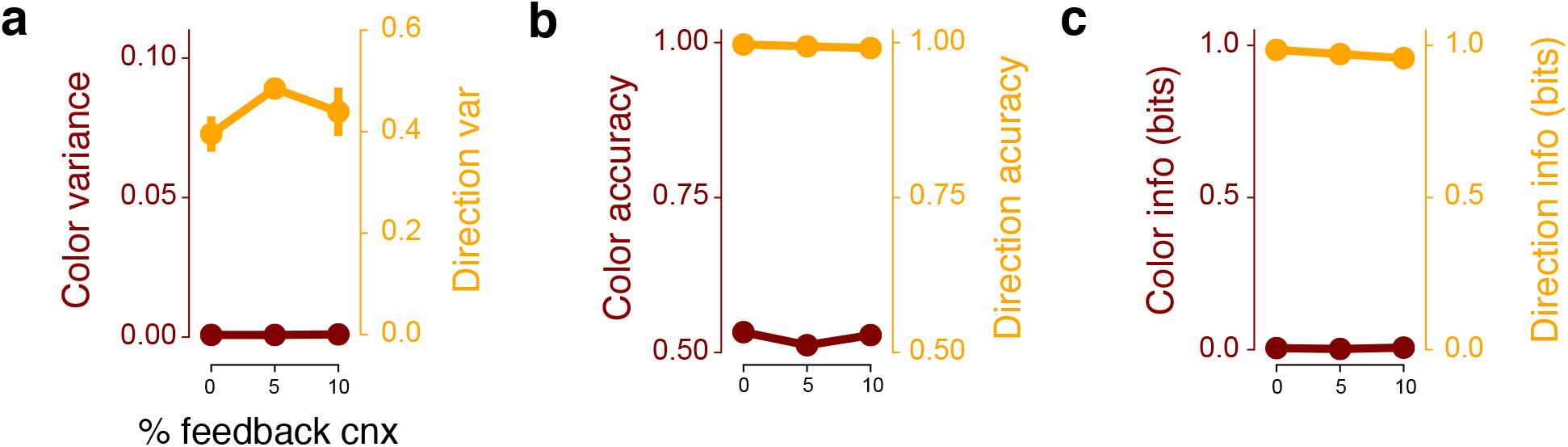
Color information is robust to varying feedback connectivity. **(a)** dPCA variance in 3-area PMd-like neurons varying the amount of feedback connectivity. RNNs exhibited nearly zero dPCA color variance across networks with 0%, 5%, and 10% feedback connections. **(b, c)** RNNs also exhibited minimal color representations, achieving nearly chance levels of decode accuracy and nearly zero mutual information.

**Figure S9:**
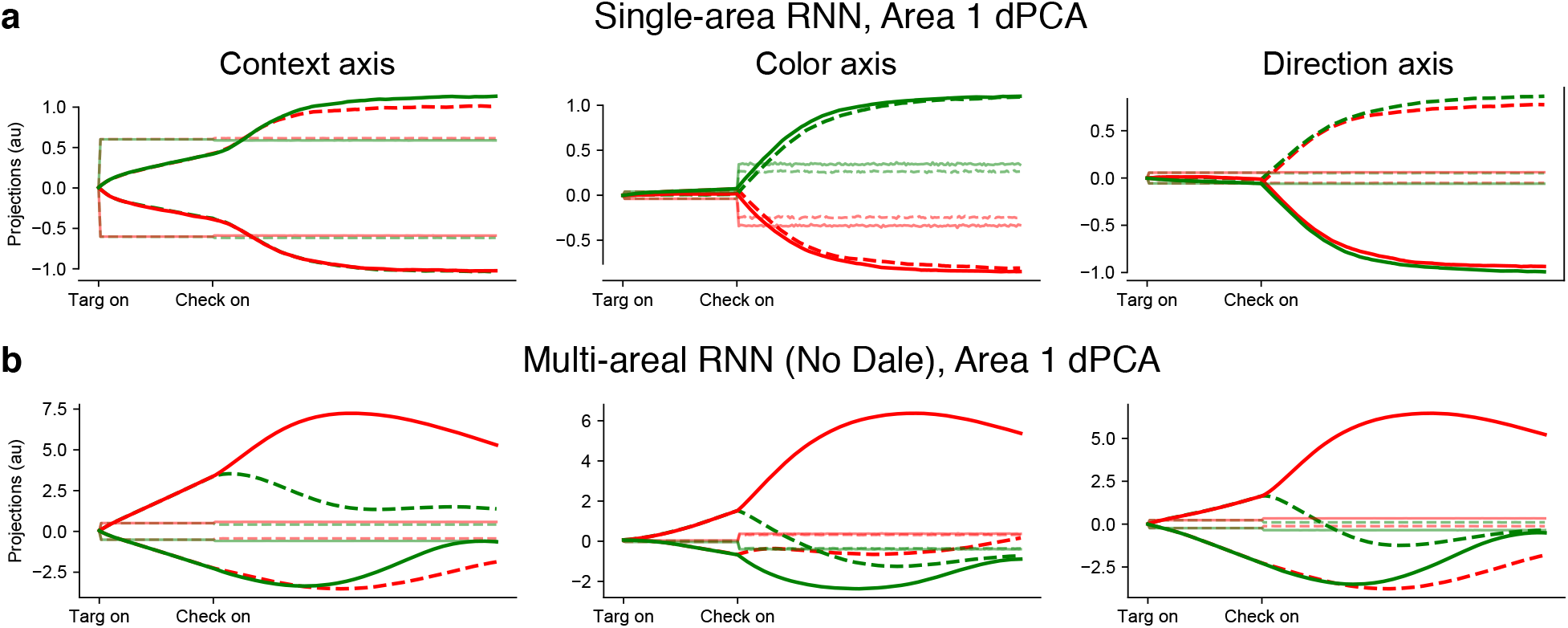
dPCA trajectories for single-area and 3-area RNNs with No Dale’s law. **(a)** Projections onto the dPCA context, color, and direction axes for a single-area RNN. dPCA was able to find axes that separate the context input, color decision, and direction decision. Importantly, in these networks, Inputs were non-zero on the direction axis. **(b)** dPCA projections for the unconstrained 3-area RNNs with color representation in Area 3. Inputs were similarly non-zero on the direction axis. The context inputs, color decision, and direction decision, had similar projection motifs.

**Figure S10:**
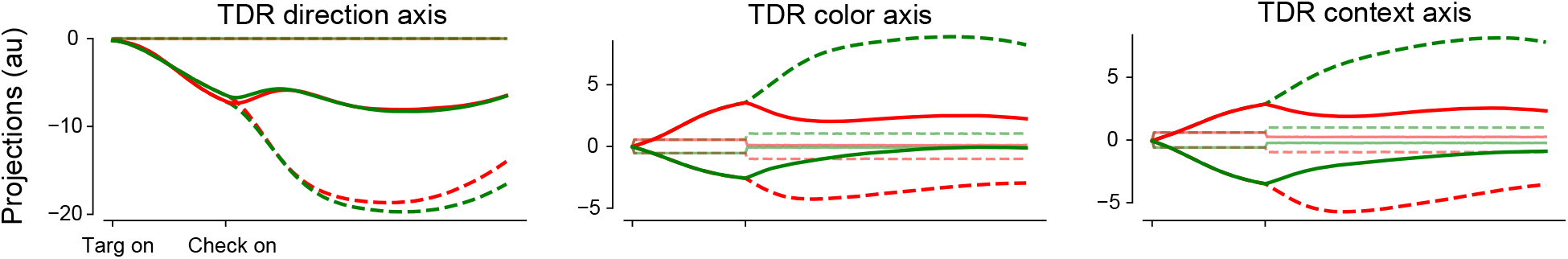
TDR results closely match dPCA results, and identifies mixed color and context axes. The direction axis separated trajectories based on the direction choice. The color and context axes had trajectory separation depending on both color and context. We did not show the orthonormalized bases, because we found that the QR decomposition was susceptible to the order in which orthonormalization was performed. This is further evidence that the color and context axes are closely aligned.

**Figure S11:**
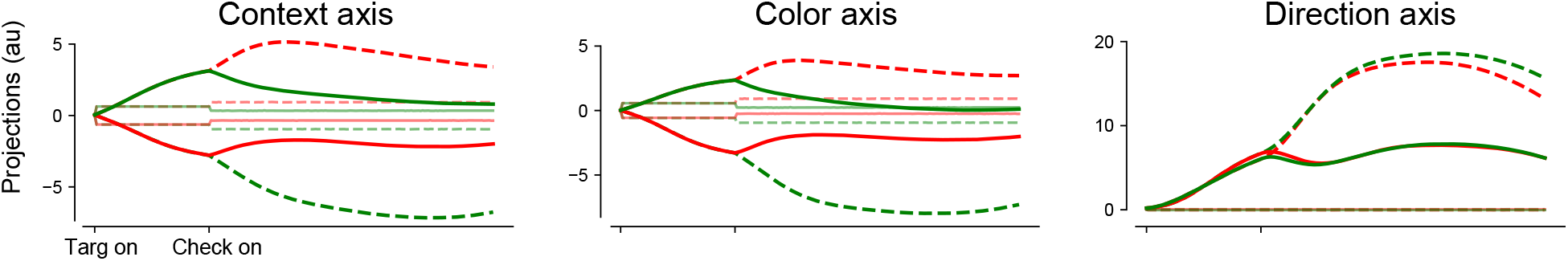
dPCA projections when only considering excitatory units. We identified the dPCA principal axes for context, color, and direction using only excitatory units. Results are consistent with the results of Fig. 4d.

**Figure S12:**
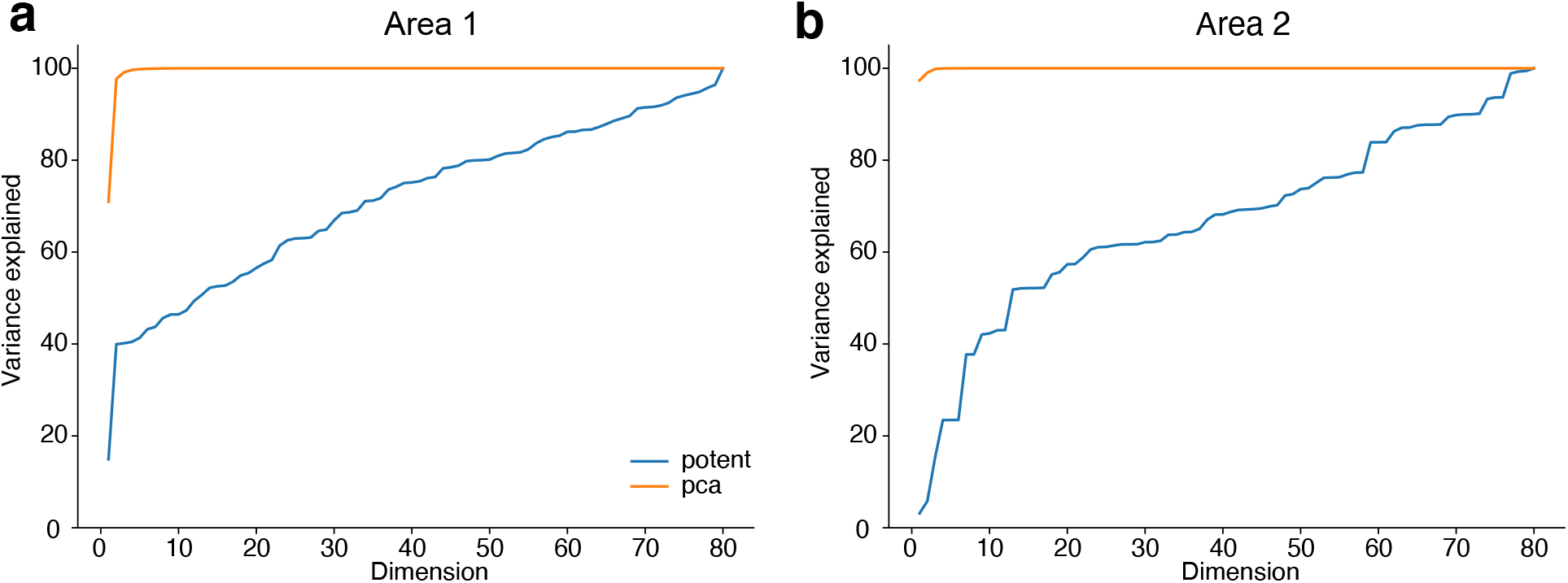
Relationship between PCs and inter-area potent space. **(a)** Variance explained of the excitatory units in Area 1 by the top principal components and top dimensions of potent space of **W**_12_, swept across all dimensions. **(b)** Variance explained of the excitatory units in Area 2 by the top principal components and top dimensions of potent space of **W**_23_, swept across all dimensions. These plots show that the connections between areas do not necessarily propagate the most dominant axes of variability in the source area to the downstream area. Excitatory units were used for the comparison because only excitatory units are read out by subsequent areas. These results were upheld when comparing to the variance explained by the top principal components obtained from all units.

**Figure S13:**
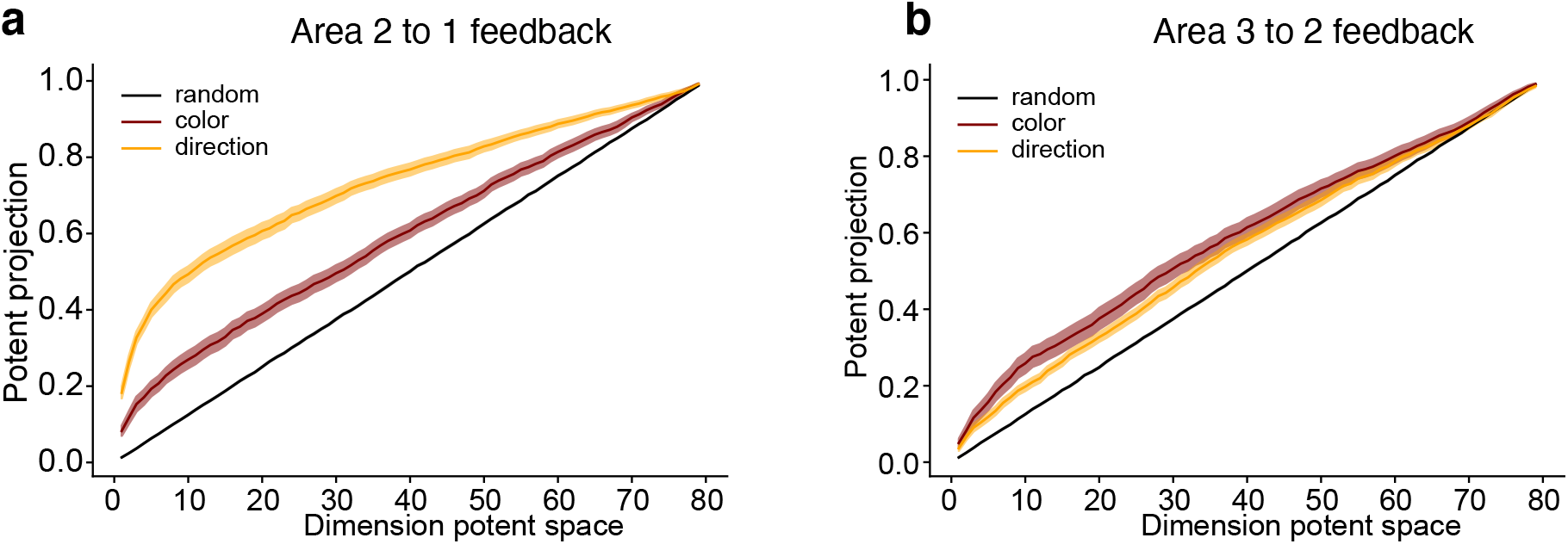
Potent projections for inter-area feedback connections between Areas 2 to 1 and Areas 3 to 2. **(a)** Alignment of the direction, color, and random axes with the inter-area feedback connections between Area 2 to Area 1. There is preferential alignment of the direction axes with the feedback connections. Color information is more aligned with feedback connections than by chance (random). **(b)** Same as **(a)**, but for Area 3 to Area 2. While the color axis is more aligned than random projections, we note that the color axis captures less than 1% variance in Area 3 (Fig 2e).

**Figure S14:**
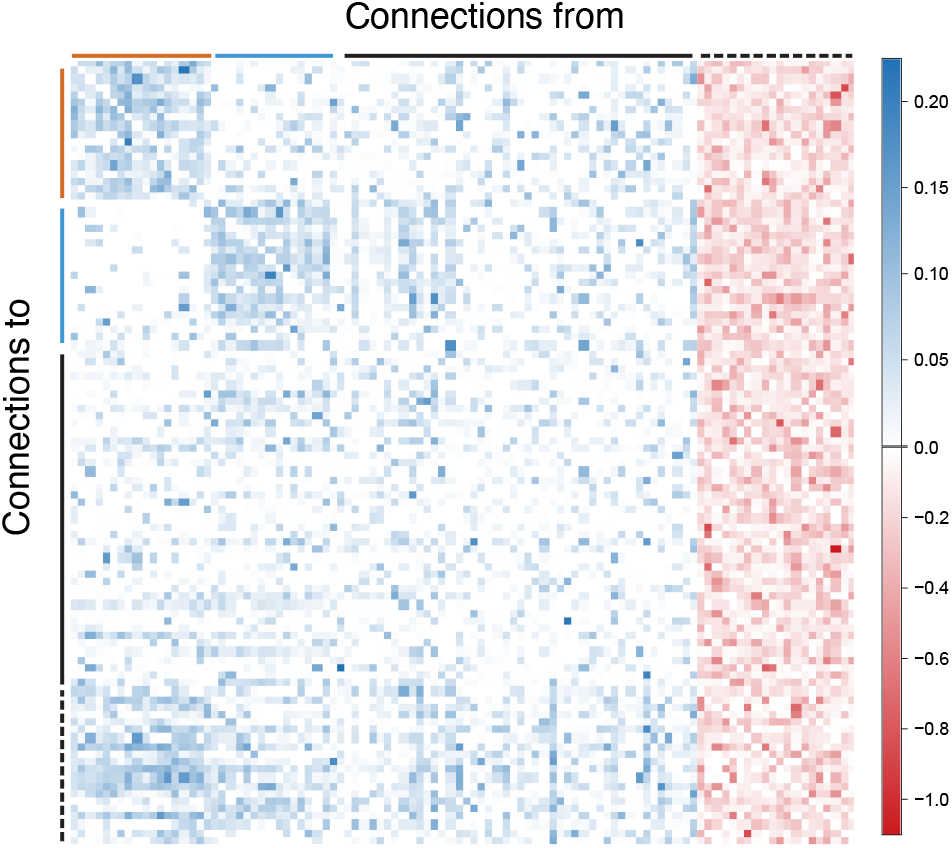
Structure of W_rec_. Full connectivity matrix of **W**_rec_, reordered so that the structured excitatory components lie at the top left. The matrix is composed of a structured excitatory component (orange and blue), a set of random excitatory units (black), and a set of inhibitory units (dashed black), with non-obvious structure. The averaged connectivity matrix is shown in Fig. 7e. We note that since we did not see clear structure in the inhibitory pool (i.e mutual inhibition), leading us to hypothesize that common inhibition led to a winner-take-all mechanism.

**Figure S15:**
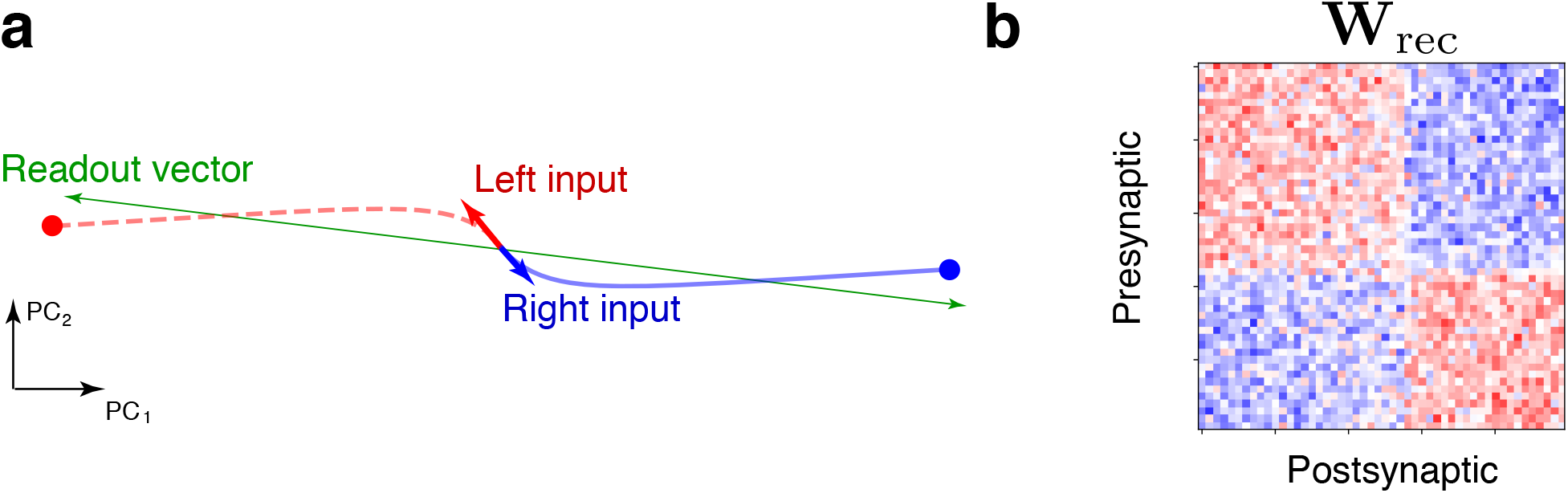
RNN trained with signed directional evidence inputs. **(a)** Projection of left (red) and right (blue) trajectories onto the top 2 PCs. The inputs are partially aligned with the readout (green). The solid dot corresponds to stable fixed points of the RNN for both input levels. **(b)** We used tensor components analysis (TCA) Williams et al. (2018) to re-sort the connectivity matrix, revealing winner-take-all structure, where two pools of neurons are self-excitatory with mutual inhibition.

### Supplementary Note 1: Viewing the CB task as an XOR task

Here we show that a nonlinearity is necessary to solve the task, proving that the task cannot be solved by the linear layer **W**_in_. First, we note that the Checkerboard task corresponds to an exclusive-or (XOR) problem. If we identify the two target configurations as 0 or 1 (corresponding to green on left, or green on right respectively, with the red target on the complement side), and the dominant checkerboard color as 0 or 1 (for green or red, respectively), then the output direction *d* (identified as 0: left, 1: right) can be seen be in Table S1.

If the representation **r** was purely input driven, then:

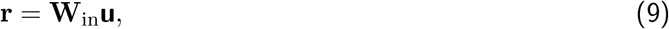

Our readout was a linear readout of the rates, i.e:

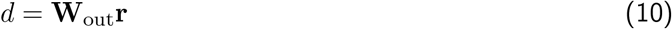

The inputs **u** are the four dimensional input we trained with. But **u** is a linear transformation of two variables: the target orientation *θ*, and checkerboard color *c*, which each can take two values. That is, if we let **q** = [*θ, c*], then, the inputs could be written as a linear transformation of **q**:

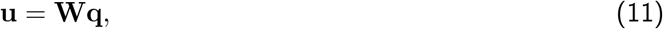

where **W** is a linear transformation. Since the mappings from **q** to *d* are all linear, they can be combined into a single linear transformation 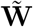, i.e.,

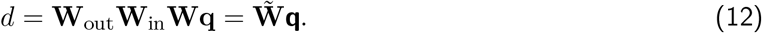

It is not possible for a linear classifier to solve the XOR problem by classifying correct outputs Goodfellow et al. (2016). Hence, the trained RNNs cannot purely be input driven, and requires nonlinearity from the recurrent interactions to solve the task. After nonlinear processing, the left or right decision could be achieved by a linear readout of the units.

**Table S1:** Checkerboard task truth table

| target configuration | color | direction |
| --- | --- | --- |
| 0 | 0 | 0 |
| 0 | 1 | 1 |
| 1 | 0 | 1 |
| 1 | 1 | 0 |

**Table S2:** Mante et al. (2013) model truth table

| context | signed color | signed motion | direction |
| --- | --- | --- | --- |
| 0 | 0 | 0 | 0 |
| 0 | 0 | 1 | 0 |
| 0 | 1 | 0 | 1 |
| 0 | 1 | 1 | 1 |
| 1 | 0 | 0 | 0 |
| 1 | 0 | 1 | 1 |
| 1 | 1 | 0 | 0 |
| 1 | 1 | 1 | 1 |

### Supplementary Note 2: Mutual Information Estimation

The entropy of a distribution is defined as

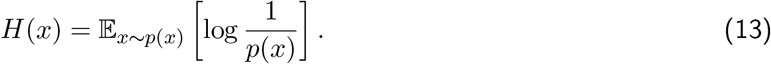

The mutual information, *I*(*X*; *Y*), can be written in terms on an entropy term and as conditional entropy term:

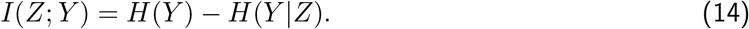

We want to show that the usable information lower bounds the mutual information:

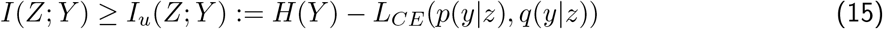

It suffices to show that:

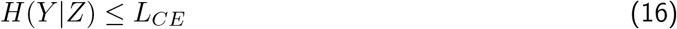

where *L_CE_* is the cross-entropy loss on the test set. For our study, *H*(*Y*) represented the known distribution of output classes, which in our case were equiprobable.

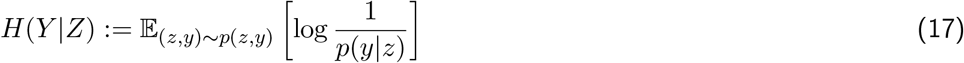

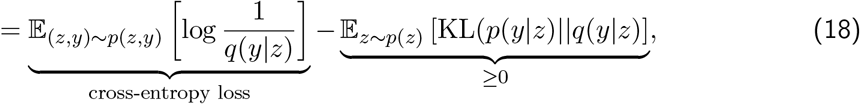

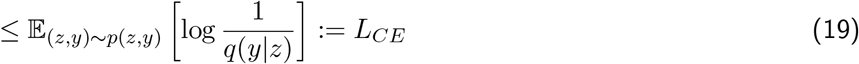

To approximate *H*(*Y*|*Z*), we first trained a neural network with cross-entropy loss to predict the output, *Y*, given the hidden activations, *Z*, learning a distribution *q*(*y*|*z*). The KL denotes the Kullback-Liebler divergence. We multiplied (and divided) by an arbitrary variational distribution, *q*(*y*|*z*), in the logarithm of equation 17, leading to equation 18. The first term in equation 18 is the cross-entropy loss commonly used for training neural networks. The second term is a KL divergence, and is therefore non-negative. In our approximator, the distribution, *q*(*y*|*x*), is parametrized by a neural network. When the distribution *q*(*y*|*z*) = *p*(*y*|*z*), our variational approximation of *H*(*Y*|*Z*), and hence approximation of *I*(*Z*; *Y*) is exact (Barber & Agakov, 2003, Poole et al., 2019).

In the paper, we additionally report the accuracy of the neural network on the test set. This differs from the cross-entropy in that the cross-entropy incorporates a weighted measure of the accuracy based on how “certain” the network is, while the accuracy does not.

## Acknowledgments

We thank Brandon McMahan for training a single-area RNN model with signed directional evidence and preparing Fig. S15. MK was supported by the National Sciences and Engineering Research Council (NSERC). CC was supported by a NIH/NINDS R00 award R00NS092972, the Moorman-Simon Interdisciplinary Career Development Professorship from Boston University, the Whitehall foundation, and the Young Investigator Award from the Brain and Behavior Research Foundation. JCK was supported by NSF CAREER 1943467, NIH DP2NS122037, the Hellman Foundation, and a UCLA Computational Medicine AWS grant. We gratefully acknowledge the support of NVIDIA Corporation with the donation of the Titan Xp GPU used for this research. We thank Laura Driscoll for helpful comments on the manuscript as well as Krishna V. Shenoy and William T. Newsome for helpful discussions on earlier versions of these results. We also thank Krishna V. Shenoy for kindly allowing us to use the data collected by Dr. Chandrasekaran when he was a postdoc in the Shenoy Lab.

## Author contributions

JCK and CC conceived of the study. MK and JCK trained RNNs and analyzed networks. MK performed the multi-area computation analyses. CC collected experimental data in PMd in Prof. Shenoy’s lab. All authors came up with analyses and conceptual frameworks to understand computation in the RNNs, discussed results, and wrote and edited the manuscript.

